# Cell types of the human retina and its organoids at single-cell resolution: developmental convergence, transcriptomic identity, and disease map

**DOI:** 10.1101/703348

**Authors:** Cameron S. Cowan, Magdalena Renner, Brigitte Gross-Scherf, David Goldblum, Martin Munz, Jacek Krol, Tamas Szikra, Panagiotis Papasaikas, Rachel Cuttat, Annick Waldt, Roland Diggelmann, Claudia P. Patino-Alvarez, Nadine Gerber-Hollbach, Sven Schuierer, Yanyan Hou, Aldin Srdanovic, Marton Balogh, Riccardo Panero, Pascal W. Hasler, Akos Kusnyerik, Arnold Szabo, Michael B. Stadler, Selim Orgül, Andreas Hierlemann, Hendrik P. N. Scholl, Guglielmo Roma, Florian Nigsch, Botond Roska

## Abstract

How closely human organoids recapitulate cell-type diversity and cell-type maturation of their target organs is not well understood. We developed human retinal organoids with multiple nuclear and synaptic layers. We sequenced the RNA of 158,844 single cells from these organoids at six developmental time points and from the periphery, fovea, pigment epithelium and choroid of light-responsive adult human retinas, and performed histochemistry. Cell types in organoids matured *in vitro* to a stable ‘developed’ state at a rate similar to human retina development *in vivo* and the transcriptomes of organoid cell types converged towards the transcriptomes of adult peripheral retinal cell types. The expression of disease-associated genes was significantly cell-type specific in adult retina and cell-type specificity was retained in organoids. We implicate unexpected cell types in diseases such as macular degeneration. This resource identifies cellular targets for studying disease mechanisms in organoids and for targeted repair in adult human retinas.

## Introduction

Human organoids are 3D cellular ensembles that are grown *in vitro* from adult or pluripotent stem cells and reproduce some morphological, functional and transcriptomic features of human organs (Clevers, 2016; Lancaster and Knoblich, 2014a). Organoids engineered to harbor disease-causing mutations or grown directly from patient cells could provide mechanistic insights into diseases.

However, human organs consist of many specialized cell types and it is not well understood how individual cell types in organoids mature, whether they converge towards the cell types of the adult organ, or how the developmental rate of organoids compares to that of the target organ. Nor is it known which disease genes retain their specificity for cell types between the target organ and its organoids or to what extent cell types in organoids have the subcellular specializations affected by disease.

The retina is a relevant model system to address these questions since its cell types have been extensively studied (Masland, 2012) and retinal organoids can be grown from human pluripotent stem cells (Meyer et al., 2011; Nakano et al., 2012; Zhong et al., 2014). Furthermore, many genes have been described that cause or contribute to vision-impairing monogenic and complex retinal diseases, such as retinitis pigmentosa and macular degeneration (Ferrari et al., 2011; Fritsche et al., 2016; Ran et al., 2014).

Retinas of humans have two distinct regions. The retinal periphery has low spatial acuity and is responsible for night-vision and different aspects of motion vision. The fovea (or macula) (Bringmann et al., 2018) is at the retinal center and drives high spatial acuity vision that is essential for reading and face recognition. Primates are the only mammals with a fovea. Retinal cells in both periphery and fovea can be divided into morphologically (Bae et al., 2018), functionally (Baden et al., 2016; Dacey et al., 2003; Roska and Werblin, 2001) and transcriptomically (Macosko et al., 2015; Peng et al., 2019; Shekhar et al., 2016; Siegert et al., 2012) different cell classes that are further divisible into cell types. The neural retina contains five layers (Dowling, 2012). Cell bodies are arranged in three distinct layers: photoreceptors in the outer nuclear layer, horizontal/bipolar/amacrine/Müller cells in the inner nuclear layer, and amacrine/ganglion cells in the ganglion cell layer. Retinal neurons make synaptic connections in two interjacent layers: the outer and inner plexiform layers. Embedded in this layered structure are astrocytes, microglial cells, and the retinal vasculature, which is composed of endothelial cells and pericytes. Conserved across vertebrates, the five-layered neural retina is covered on the photoreceptor side by the retinal pigment epithelium and by the choroid, which contains endothelial cells, pericytes, fibroblasts, and melanocytes (Nickla and Wallman, 2010). We refer to the neural retina, the retinal pigment epithelium and the choroid together as ‘the retina’. Cells in the retina display a number of functionally important subcellular specializations. In photoreceptors, for example, the outer segment captures light, the connecting cilium enables the transport of outer-segment specific molecules, the inner segment hosts mitochondria that produce energy, and the ribbon synapse permits graded signal transmission of sensory information. The distinguishing characteristics of the cell types and their subcellular specializations are associated with the expression of specific genes that are frequently implicated in retinal disease (Ran et al., 2014). The transcriptomes of some adult human retinal cell types have been described (Kim et al., 2019; Peng et al., 2019); however, the cells were sampled from donor retinas post mortem after six hours in a hypoxic state. Hypoxia leads to irreversible damage to the retina within 20 minutes (Osborne et al., 2004). The transcriptomes of cell types in human adult retinal pigment epithelium and choroid have not yet been described.

Human retinal organoids have been used to study retinal development and disease (Foltz and Clegg, 2019), but currently used organoids are not five-layered like the real human retina. They are also difficult to produce in large quantities because their generation often involves a microdissection step (Zhong et al., 2014). An additional barrier to modeling genetic diseases of the retina is the lack of a comprehensive quantitative comparison of gene expression between cell types in organoids and human retinas (Collin et al., 2019; Kim et al., 2019). For cell types in retinal organoids it is therefore not well understood whether and when their gene expression matches that of the adult human retina. For many retinal disease genes, it is not known in which cell types of the adult human retina they are expressed or at what age organoid cell types reproduce this expression.

Here, we report the development of retinal organoids with three nuclear and two synaptic layers from human induced pluripotent stem cells (iPSCs) and their production in large quantities. We obtained single-cell transcriptomes from 62,136 cells dissociated from these developing human multilayered organoids at six different time-points spanning the 38 weeks of human gestation. Deep learning analysis of these transcriptomes revealed progressive maturation of retinal cell classes and showed that organoid transcriptomes reached a stable, developed state by week 30. The rate of transcriptomic changes during organoid development was similar to the developing human retina *in vivo*. We then developed a procedure to obtain adult human retinas that were exposed to less than five minutes of hypoxia and maintained light responses and functional retinal circuits for 16 hours *ex vivo*. We sequenced RNA from 96,708 single cells from the peripheral and foveal retina, including the pigment epithelium and choroid. Comparing periphery to fovea, we identified regional characteristics of cell types and, by comparing organoid to organ, we showed that transcriptomes of organoid cell types converge to those of adult human peripheral retinas. In the context of cell types, we also compared developed organoids and adult human retinas in their expression of genes associated with retinal diseases. The resulting genetic disease maps showed retinal diseases to be cell-type specific and that cell-type specificity is preserved in organoids. The resources we describe here provide an iPSC line and a high-throughput method to build retinal organoids, with reproducible organ features and transcriptomes, for modelling retinal disease. Furthermore, we provide a comparative atlas of cell-type transcriptomes of human retinal organoid and healthy adult human retina that allows identification of cellular targets for studying disease mechanisms in organoids and targeted repair in adult human retinas.

## Results

### Multilayered human retinal organoids produced in quantity

We aimed to develop human retinal organoids with three nuclear and two synaptic layers from iPSCs, building on the observation that iPSC lines vary in their potential to produce retinal organoids: some gave rise to organoids while others did not. As organoid differentiation is a nonlinear process (Dahl-Jensen and Grapin-Botton, 2017), the qualitatively different outcomes could be due to differences in the initial condition of the iPSCs, namely their genetic origin or their epigenetic and transcriptomic states at the time of differentiation (Kilpinen et al., 2017; Ortmann and Vallier, 2017). Therefore, we screened 21 iPSC lines for the formation of multilayered retinal organoids. Eight of the lines formed retinal organoids and six lines had a layered appearance under the light microscope that persisted for more than 100 days in culture (Supplemental Table 1). We visualized the nuclear layers in organoid sections using a Hoechst dye and the synaptic layers using an antibody against Bassoon. The 01F49i-N-B7 (short name: F49B7) line formed retinal organoids with three nuclear and two synaptic layers, matching the cellular organization of the adult retina (Figure 1). F49B7 iPSCs expressed pluripotency markers, could be differentiated into all the three germ layers, and had a normal karyotype (Supplemental Figure 1).

**Figure 1.**
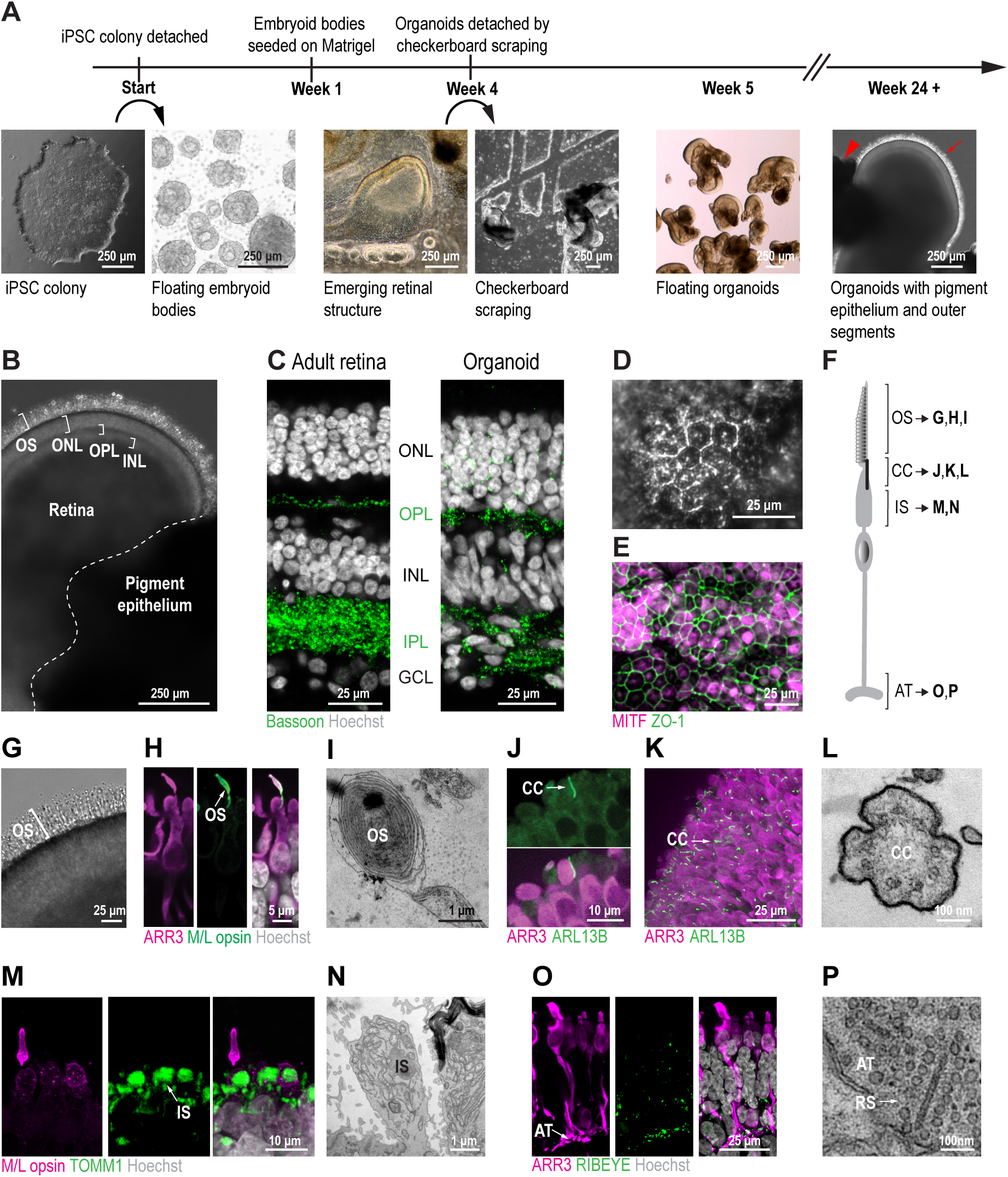
Multilayered human retinal organoids produced in quantity. (A) Timeline of the organoid protocol with example brightfield images at important stages. Red arrowhead, pigment epithelium; red arrow, outer segments. (B) Brightfield image of an organoid. OS, outer segment; ONL, outer nuclear layer; OPL, outer plexiform layer; INL, inner nuclear layer. (C) Confocal images. Left, adult retina. Right, organoid. Green, Bassoon antibody (synaptic marker); white, Hoechst (nucleus marker). ONL, outer nuclear layer; OPL, outer plexiform layer; INL, inner nuclear layer; IPL, inner plexiform layer; GCL, ganglion cell layer. (D) Brightfield image of organoid pigment epithelial cells with black pigmentation. (E) Confocal image of pigment epithelial cells (maximum intensity projection). Magenta, MITF antibody; green, ZO-1 antibody (pigment epithelial cell markers). (F) Illustration of photoreceptor subcellular compartments. OS, outer segment; CC, connecting cilium; IS, inner segment; AT, axon terminal. (G – P) Organoid photoreceptors. (G – I) Outer segment. (G) Brightfield image. (H) Confocal images. Magenta, ARR3 antibody (cone marker); green, M/L opsin antibody (cone outer segment marker); white, Hoechst (nucleus marker). (I) Electron microscope image. Diagonal section of membrane discs. (J – L) Connecting cilium. (J – K) Confocal images. Magenta, ARR3 antibody; green, ARL13B antibody (cilium marker). (K) Maximum intensity projection showing organoid surface. (L) Electron microscope image. (M – N) Inner segment. (M) Confocal image. Magenta, M/L opsin antibody; green, TOMM1 antibody (mitochondrium marker). (N) Electron microscope image. (O – P) Axon terminal. (O) Confocal image. Magenta, ARR3 antibody; green, RIBEYE antibody (ribbon synapse marker). (P) Electron microscope image. AT, axon terminal; RS, ribbon synapse.

Current organoid production methods include a manual microdissection step that precludes the fabrication of organoids in large quantities. Therefore, we developed a method that omits microdissection (Figure 1A) and characterized it using the F49B7 line. Instead of microdissecting optic-cup-like structures that developed after three weeks of retinal 2D culture on Matrigel-coated plates, we dislodged the contents of the entire plate by scraping along a checkerboard pattern. Scraping yielded numerous separate tissue pieces that developed into retinal organoids, characterized by phase-bright neuroepithelium (Supplemental Figure 2). The procedure is faster than manual dissection (five minutes vs. 40 minutes per plate) and yielded retinal organoids with a stable (> eight months) layered appearance (Figure 1B – C) in quantities 20 times higher than the microdissection method resulting in 21.7 ± 10.3 retinal organoids per well of a 6-well plate (mean ± SD, *n* = 108 organoids, three experiments, at week 38). Thus, the overall throughput of the checkerboard method was at least 160 times higher than the manual microdissection method. Of the organoids produced by the checkerboard method, 76% were retinal organoids. The remaining products were either pigment epithelium spheroids (8.5%) or undefined structures (15.5%). Of the retinal organoids, 97% contained isolated patches of pigment epithelium at week 38 (*n* = 135 organoids, three experiments). The pigment epithelium was composed of characteristic pigmented hexagonal cells (Figure 1D) expressing the pigment epithelium markers MITF and ZO-1 (Figure 1E).

We then asked whether organoid photoreceptors contain key subcellular structures (Figure 1F). The photoreceptors in 97% of organoids grew processes resembling outer segments (Figure 1G), which at week 32 had a length of 47 ± 7.5 μm (mean ± SD, *n* = 31 organoids, four experiments). Consistent with the notion of outer segments, cone opsin (Figure 1H) and rhodopsin (Supplemental Figure 2) were both localized to the protruding processes and we observed regularly spaced membrane discs (Figure 1I; Supplemental Figure 2). In addition, organoid photoreceptors had connecting cilia (Figure 1J – L; Supplemental Figure 2), inner segments containing mitochondria (Figure 1M – N; Supplemental Figure 2), and axon terminals containing ribbon synapses (Figure 1O – P).

### Organoids stabilize to a developed state

We approached the question of when organoids in culture are ‘developed’ by performing single-cell RNA sequencing at six time points during organoid development: 6, 12, 18, 24, 30 and 38 weeks (Supplemental Figure 2). We analyzed 62,136 cells from F49B7 organoids using the 10x Genomics Chromium platform (*n* = 12 individual organoids, *n* = 7 pooled organoids) (Figure 2A). We embedded the transcriptomes of organoid cells, pooled across all time points, within a 2D map using the deep learning-based scVis algorithm (Ding et al., 2018) (Figure 2B). On the scVis map, each point represents the transcriptome of a single cell and adjacent points correspond to transcriptomes that are similar to each other.

**Figure 2.**
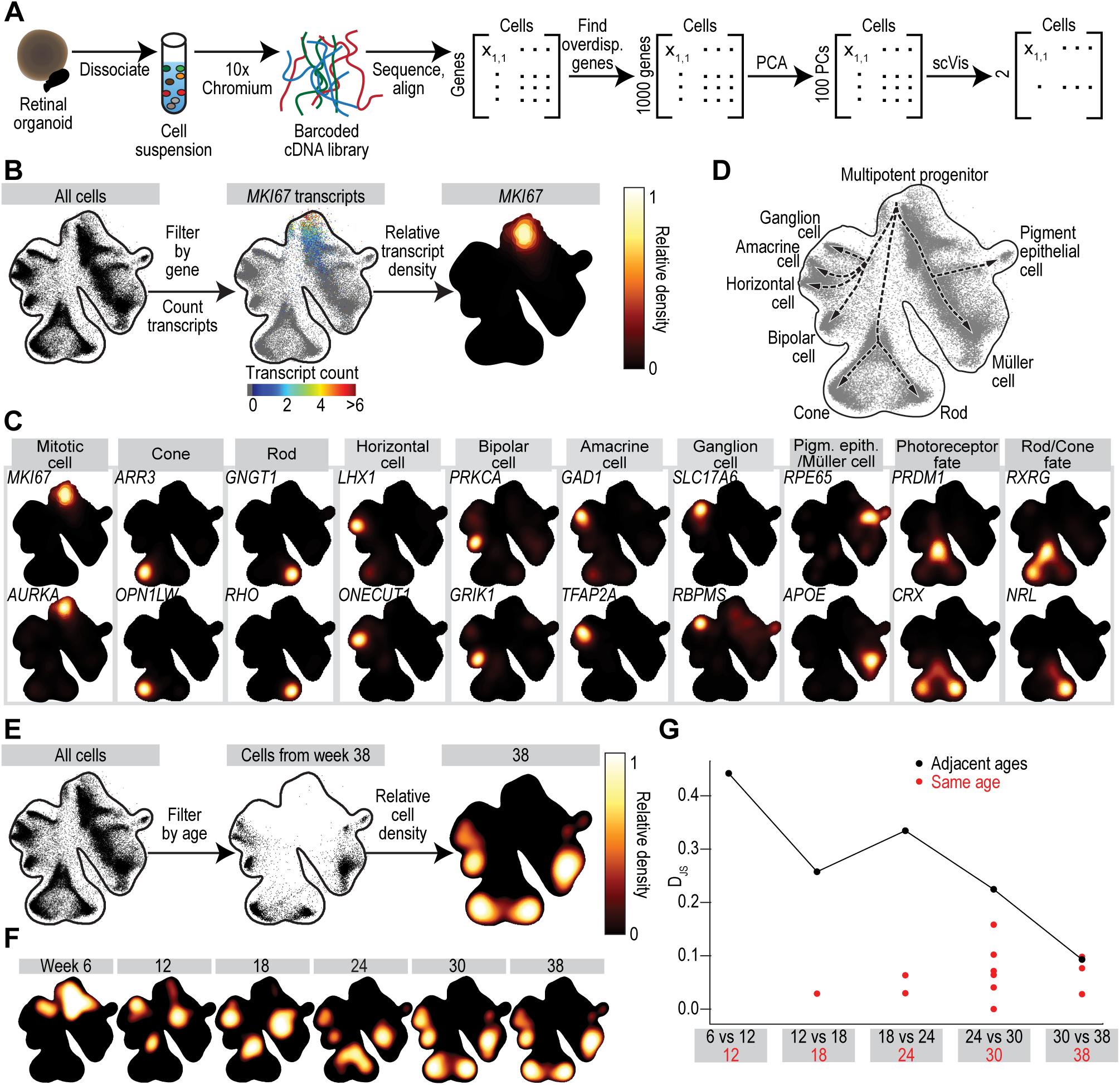
Organoid transcriptomes reach a stable state. (A) Experimental and bioinformatic steps from organoids to single-cell transcriptomes visualized in scVis maps. PCA, principal component analysis; PC, principal component. (B) Left, scVis map of cells across all ages (weeks 6 – 38). Each point represents the transcriptome of a single cell. Black line, isodensity contour. Middle, scVis map with transcript counts of a single gene (here, mitosis marker *MKI67)* color-coded; colormap at bottom. Right, heatmap of relative transcript density for *MKI67*; colormap at right. (C) Heatmap of relative transcript density for genes marking mitotic cells, retinal cell classes, and retinal precursors. (D) Cell classes marked on scVis map according to C. Arrows, inferred developmental trajectories. (E) Left, scVis map of cells across all ages. Middle, a black point represents a cell from an organoid of a specific age (here week 38). Right, heatmap of relative cell density at week 38; colormap at right. (F) Heatmap of relative cell density at six different ages. (G) Black line, Jensen-Shannon divergence (*D*_JS_) of organoid transcriptomes at adjacent ages. Red dots, comparison of individual organoids from the same age and batch.

Within the scVis map, single-cell transcriptomes were distributed along a leaf-like manifold. We located on this manifold the transcriptomes of cells expressing known genetic markers of retinal progenitors or specific retinal cell classes (Figure 2B – D). Mitotic cell division markers *MKI67* and *AURKA,* expressed in retinal progenitor cells, identified transcriptomes at the top corner of the manifold (Figure 2C). Two pools of progenitor cells were present, in one the neurogenic marker *NEUROD1* was upregulated (60× upregulated, *P* = 1.8×10^-308^), and in the other the gliogenic marker *PAX2* was upregulated (37× upregulated, *P* = 5.8×10^-12^) (Supplemental Figure 3). Markers for cones (*ARR3*, *OPN1LW*), rods (*GNGT1*, *RHO*), horizontal cells (*LHX1*, *ONECUT1*), bipolar cells (*PRKCA*, *GRIK1*), amacrine cells (*GAD1*, *TFAP2A*), ganglion cells (*SLC17A6*, *RBPMS*), pigment epithelial cells (*RPE65*) and Müller cells (*APOE*) each identified transcriptomes at different distal tips of the manifold. Bipotent photoreceptor precursors, identified by the fate markers *PRDM1* and *CRX*, were located inward of rods and cones. The genes defining rod (*NRL*) or cone (*RXRG*) fate commitment were detected in transcriptomes distributed along a path between the transcriptomes of photoreceptor precursors and the corresponding mature cluster. Thus, an ordered series of cell transcriptomes across the leaf-like manifold captured a progression of gene expression patterns reached by cells as they develop from multipotent progenitors to differentiated retinal cell classes (Figure 2D).

We next investigated the transcriptomes of cells at different ages within the manifold and how their locations change with time (Figure 2E – F). The 6-week-old cell transcriptomes were concentrated in the manifold’s upper region and at each subsequent time point the distribution shifted closer to the regions containing transcriptomes of differentiated cell classes. The temporal order of appearance of neural retinal cells was: ganglion cells, photoreceptor precursors, horizontal cells, amacrine cells, bipolar cells, and Müller cells. Although ganglion cells were abundant at 12 – 18 weeks, they were rare by week 24: ganglion cell marker *SLC17A6* was detected in 0.36% of organoid cells between weeks 24 and 38. To quantify changes in gene expression within the 6-to 38-week time window of organoid development, we calculated the Jensen-Shannon divergence (*D*_JS_) of distributions shown on scVis maps between organoids at the same or different ages (Supplemental Figure 3). *D*_JS_ provides a measure of dissimilarity between distributions, which takes a value of zero for perfectly overlapping distributions and one for non-overlapping distributions. The mean *D*_JS_ between individual organoids from the same batch and age stayed below 0.1 through week 30 (*D*_JS_ = 0.073 ± 0.040, mean ± SD) and 38 (*D*_JS_ = 0.068 ± 0.030) (Figure 2G). The curve of *D*_JS_ plotted between adjacent ages as a function of age trended downward with increasing age (Figure 2G), reaching its minimum *D*_JS_ = 0.093 between weeks 30 and 38, a value that was within one standard deviation of the *D*_JS_ values for transcriptomes observed at the same age. Thus, the gene expression patterns of organoids stabilize by week 30 and at the same age the transcriptomes of organoids are reproducible. On this basis, we refer to organoids at the 30- and 38-week time points as ‘developed organoids’.

### Matching rates of organoid and retina development

The rate of development of an organoid *in vitro* may differ from that of the retina *in vivo*. To compare the two rates, we first cross-correlated the bulk transcriptomes of organoids, obtained by combining single-cell transcriptomes, to published bulk transcriptomes of human retina between seven and 20 weeks in developmental age (Hoshino et al., 2017). The cross-correlation matrix between organoid and retina had a strong diagonal trend (Figure 3A), with some correlations along the diagonal reaching 0.8, suggestive of comodulation in gene expression during development. To quantify developmental transcriptomic changes, we generated two self-to-self correlation matrices, one for retina (Figure 3B) and one for organoids (Figure 3C). In these matrices, gene correlations peaked for adjacent sample ages and extended for many weeks in either direction suggesting that it is possible to predict the retina-equivalent age of developing organoids. We therefore created a linear model that was trained to predict retinal age using the retinal transcriptomes (Figure 3D). The model was validated on the transcriptomes of the developing retina (Figure 3E). The trained model was then applied to the organoid transcriptomes to predict their retina-equivalent developmental age. Remarkably, when the model-predicted, retina-equivalent developmental ages of the organoids were plotted against their actual developmental ages, the data points were close to the unity line (Figure 3E). Thus, the rates of development of organoids *in vitro* and retina *in vivo* are similar.

**Figure 3.**
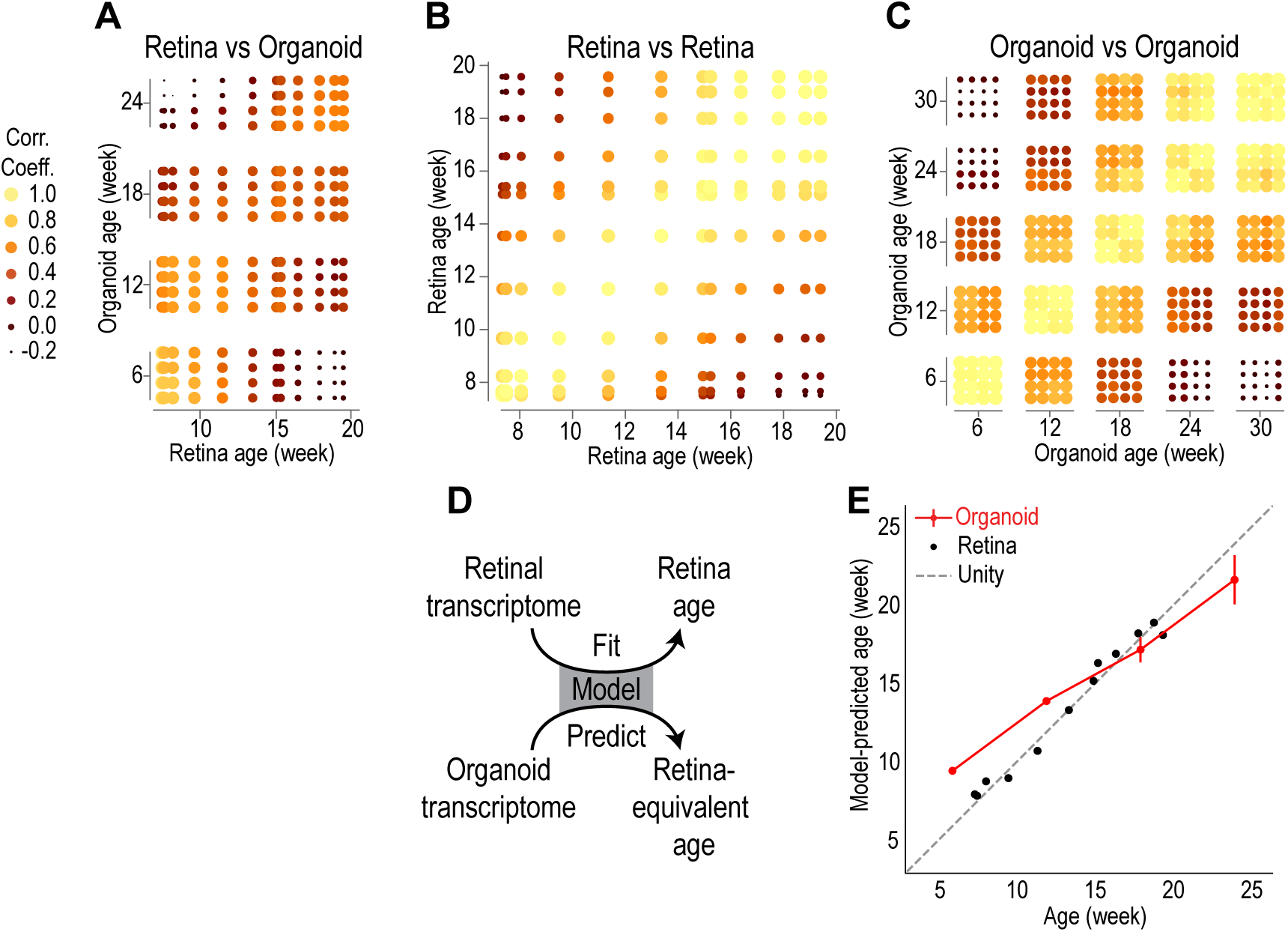
Rate of organoid development *in vitro* matches that of the retina *in vivo*. (A) Correlation in gene expression between developing organoid and developing retina. Each point is a correlation coefficient between two samples. Point size and color, correlation strength; scale at left. (B) Correlation between retinas at different ages. (C) Correlation between organoids at different ages. (D) A model, trained to predict retina age based on retinal transcriptome data, is applied to predict retina-equivalent age of organoid samples. (E) Red line, model-predicted retina-equivalent age of organoids versus their age in culture (mean ± three SE). Black points, model-predicted age of retinas versus their developmental age. Dashed grey line, one-to-one correspondence between age and model-predicted age.

### Cell-type transcriptomes in functional adult human retinas and developed organoids

Organoids have distinct and stable transcriptomically defined cell classes by week 30. To determine how close the transcriptomes of these cell classes are to those of the adult retina, we sequenced RNA of single cells from retinas of human adult multi organ donors. Since a post mortem retina is irreversibly damaged after 20 minutes in hypoxia (Osborne et al., 2004), we combined rapid surgery and eye-cup preparation with an oxygenation and perfusion-providing transportation device to reduce the ischemic window experienced by donor retinas to less than five minutes (Figure 4A). We tested the functional state of the peripheral neural retina and, in parallel, used neural retina, retinal pigment epithelium, and choroid from the periphery and fovea for single-cell RNA sequencing.

**Figure 4.**
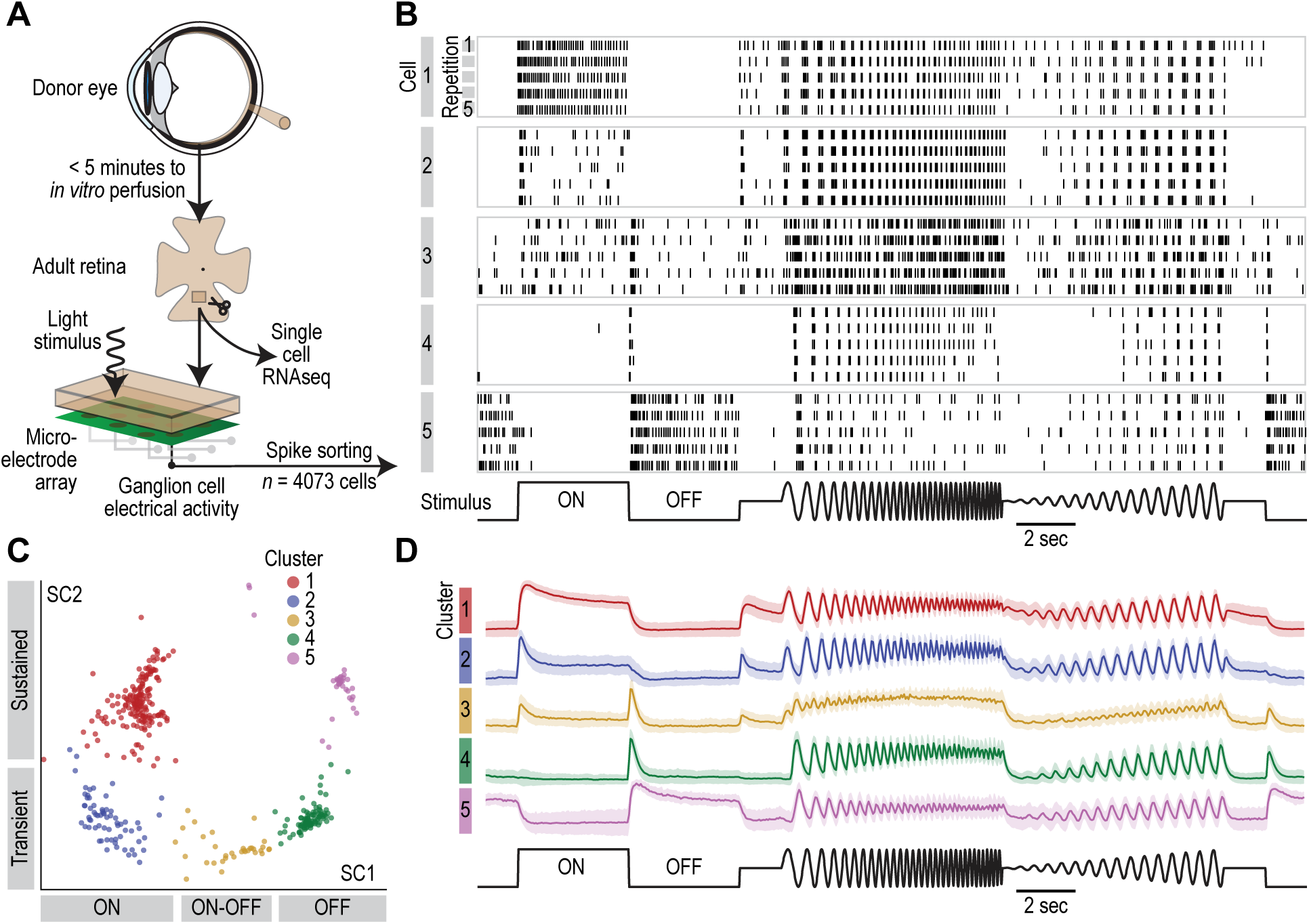
Post mortem adult retina with light-responses. (A) Scheme of the procedure to obtain adult retinas for single-cell RNA sequencing and electrophysiology. (B) Vertical lines, action potentials fired by five example ganglion cells (horizontal boxes) during five repetitions (rows within boxes) of a light stimulus (at bottom). (C) scVis map of light responses from a sample of ganglion cells at the same retinal location; each point represents the firing rate characteristics of an individual cell in response to visual stimulation. Colors, Infomap clusters. Labels at the two axes, response characteristics of cells in that region of the scVis map. (D) Colored lines, normalized firing rate of cells within each cluster during the stimulus. Shaded regions, ±1 SD. Colors, Infomap clusters as in C.

We assessed the state of the peripheral neural retina by quantifying different aspects of light-induced activity, using a high-density microelectrode array to record ganglion cells that fired action potentials during visual stimulation. Remarkably, 82% of the recorded ganglion cells (3550 cells of 4073) showed significant light responses (*P* < 0.01; Pearson correlation) to a series of spatially uniform steps, frequency chirps, and intensity sweeps (Figure 4B). Of the responding cells, 50% had a peak firing rate above 775 Hz and 95% above 52 Hz (Supplemental Figure 4). Using unsupervised Infomap clustering (Rosvall and Bergstrom, 2008), we isolated at least five distinct response clusters (Figure 4C). Cells of the first and second cluster showed responses (Figure 4D) at light onset (ON cells), with an increase in firing rate that was either sustained (cluster 1) or transient (cluster 2). The third cluster had ON-OFF cells that were transient, spiking briefly at both light onset and offset. The responses of the fourth and fifth clusters were the converse of clusters one and two, spiking at light offset (OFF cells) in either a transient (cluster four) or sustained (cluster five) manner. The observed variety of ganglion cell light responses is consistent with an adult retina that is healthy at multiple levels in its circuitry (Field and Chichilnisky, 2007): the ON and OFF responses show that the split of information from photoreceptors to ON and OFF bipolar cells is intact, transient responses show functional feedback and feedforward inhibition from horizontal and amacrine cells, and the high firing rates show efficient light capture and transmission of information across retinal synapses.

From these light responsive adult retinas, we compared the transcriptomes of individual peripheral and foveal cells (total of 96,708 cells; *n* = 74,558 peripheral retinal cells from *n* = 6 eyes; *n* = 22,150 foveal retinal cells from *n* = 5 eyes; from 3 donors) with those of cells from developed organoids (*n* = 25,143 cells from *n* = 8 individual organoids). The average number of unique transcripts per cell was 2,508 in the peripheral retina, 7,146 in the foveal retina, and 4,305 in developed organoids. We identified transcriptome-based cell types by Infomap clustering of single-cell transcriptomes in the peripheral retina, foveal retina and developed organoids independently (Figure 5A). We quantified the quality of clustering by cluster purity and by cluster stability (Shekhar et al., 2016). Cluster purity and stability were both high in the peripheral retina, foveal retina, and in developed organoids (peripheral retina: mean / 5^th^ percentile purity = 0.97 / 0.87, stability = 0.96 / 0.88; foveal retina: purity = 0.96 / 0.78, stability = 0.95 / 0.81; developed organoids: purity = 0.94 / 0.79, stability = 0.93 / 0.80) (Supplemental Figure 5). We visualized gene expression of cells using scVis embedding and represented Infomap clusters on the scVis map by different colors (Figure 5B – E). We chose scVis because it preserves distances between clusters.

**Figure 5.**
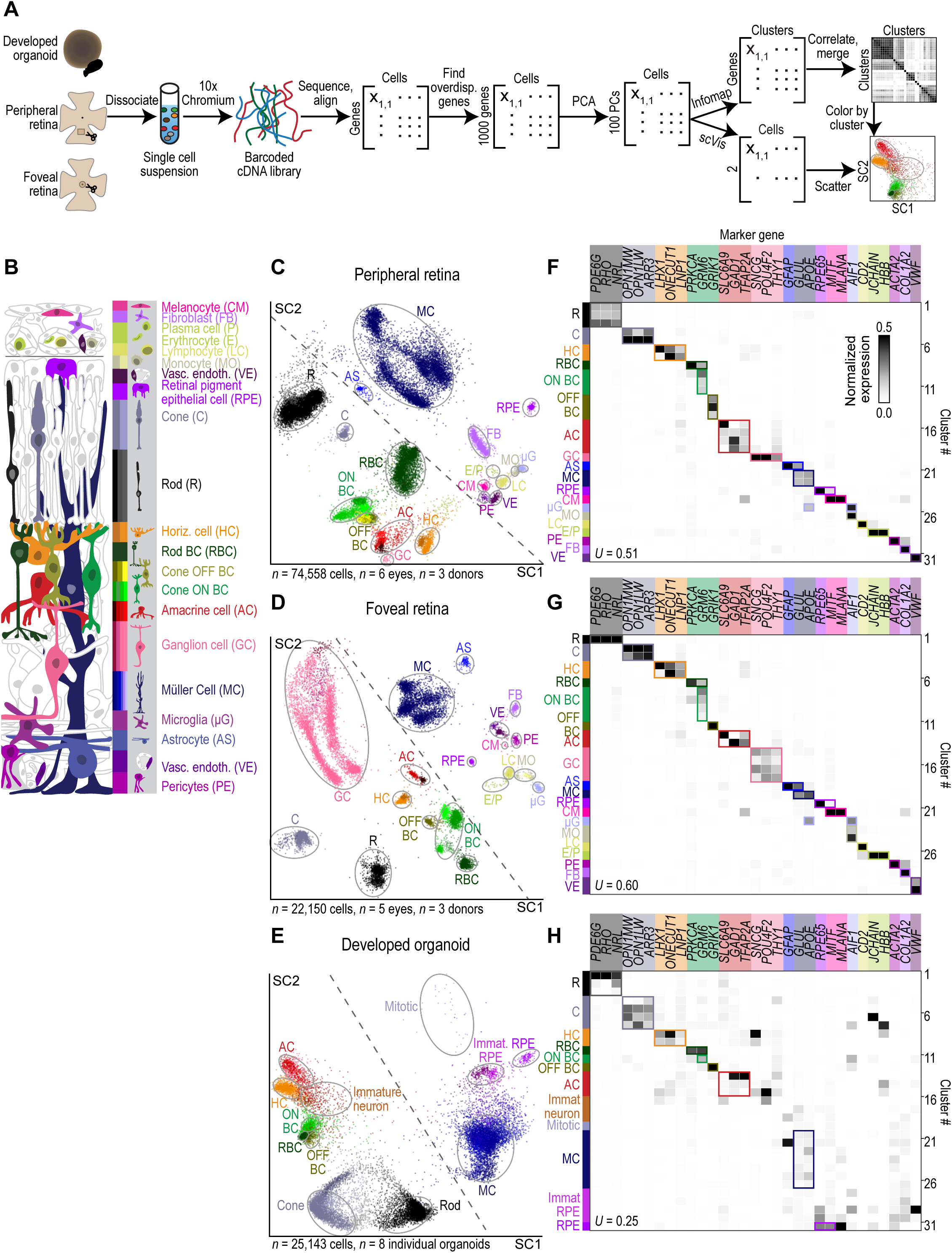
Cell types of the adult human retina and its organoids. (A) Experimental and analytical workflow, from starting tissue to an scVis map with points colored according to Infomap clusters. (B) Illustration, retinal cell classes and types. Color bar, selected cell types and classes; shades of the same color, cell types within a cell class. PCA, principal component analysis; PC, principal component. (C – E) scVis maps of transcriptomic cell types within (C) peripheral retina, (D) foveal retina and (E) developed organoids (weeks 30 and 38). Colored/circled/labeled regions, cell classes and types as in B. Dashed lines, approximate division between neuronal and non-neuronal cell types within the scVis map. (F – H) Normalized expression of known marker genes (columns) within Infomap clusters (rows) for (F) peripheral retina, (G) foveal retina and (H) developed organoids. Colormap at top right of F, intra-gene normalized expression. Color of boxes around expression blocks, cell class type identity of Infomap clusters. *U*, uncertainty coefficient.

We detected 31 transcriptome-based cell types (Infomap clusters) in peripheral retina (Figure 5C, F), 30 in the foveal retina (Figure 5D, G), and 31 in developed organoids (Figure 5E, H). These transcriptome-based cell types were compared to known marker-defined cell types or classes (Figure 5F – H) using the uncertainty coefficient (*U*). When *U* takes a value of one, the marker-defined cell types explain all of the variability in the transcriptome-based cell types; when *U* takes a value of zero, none of the variability is explained. Significance was assigned to an uncertainty coefficient by comparing it to uncertainty coefficients generated by shuffling cluster labels. The highly significant uncertainty coefficients (peripheral retina: *U* = 0.51, *P* = 3.4×10^-16^; foveal retina: *U* = 0.60, *P* = 6.8×10^-17^; developed organoid: *U* = 0.25, *P* = 1.9×10^-13^) suggest correspondence between the transcriptome-based cell types and known marker-defined cell types. We then investigated the topology of transcriptome-based cell types on scVis maps. Neuronal and non-neuronal cell types were arranged on opposite sides of the maps and comprised several clusters of cells that also showed functionally relevant grouping. For example, rod bipolar cells, which are a type of ON bipolar cells, were adjacent to ON cone bipolar cells, which were themselves adjacent to OFF cone bipolar cells, which were in turn adjacent to other interneurons. This topology was conserved in all three scVis maps − peripheral retina, foveal retina, and developed organoids. Therefore, the distance between two cell clusters on scVis maps in some cases reflects similarity in function between the cell clusters.

In the peripheral retina, we detected all known neuronal cell classes as well as many subclasses and distinct cell types, such as rods and cones, two types of horizontal cells, seven types of bipolar cells in three major groups (rod bipolar, ON cone bipolar and OFF cone bipolar), four major subclasses of amacrine cells (one GABAergic, one glycinergic and two with a mix of both), and ganglion cells. Of the neuronal cells in the foveal retina, we detected rods and cones, two types of horizontal cells, six types of bipolar cells, GABAergic and glycinergic amacrine cells, and four types of ganglion cells. The following non-neuronal cells were detected in both peripheral and foveal retina: Müller cells, astrocytes, microglia, monocytes, lymphocytes, choroidal melanocytes, pigment epithelial cells, fibroblasts, pericytes, and vascular endothelial cells.

In developed organoids, major retinal neuronal cell classes and some cell types were present, including rods and cones, horizontal cells, ON and OFF bipolar cells, and amacrine cells. The only non-neuronal cell types detected were Müller cells and pigment epithelial cells. Regions of the scVis embedding associated with cells from younger organoids included 8.85% of organoid cells, indicating some cells had not finished developing.

The ratios of different cell classes were significantly different (χ^2^ test; *P* < 10^-8^) between the peripheral and the foveal retina (Figure 6A). Rods were more frequent in the peripheral retina (rods: periphery / fovea, 60.1% / 10.0%) while cones, horizontal, bipolar, amacrine, ganglion and glial cells were less numerous in the peripheral retina than the foveal retina (cones: periphery / fovea, 2.7% / 7.2%; horizontal cells: 1.6% / 5.4%; bipolar cells: 13.4% / 18.3%; amacrine cells: 0.1% / 0.3%; ganglion cells: 0.2% / 31.3%; glial cells: 19.6% / 20.5%).

**Figure 6.**
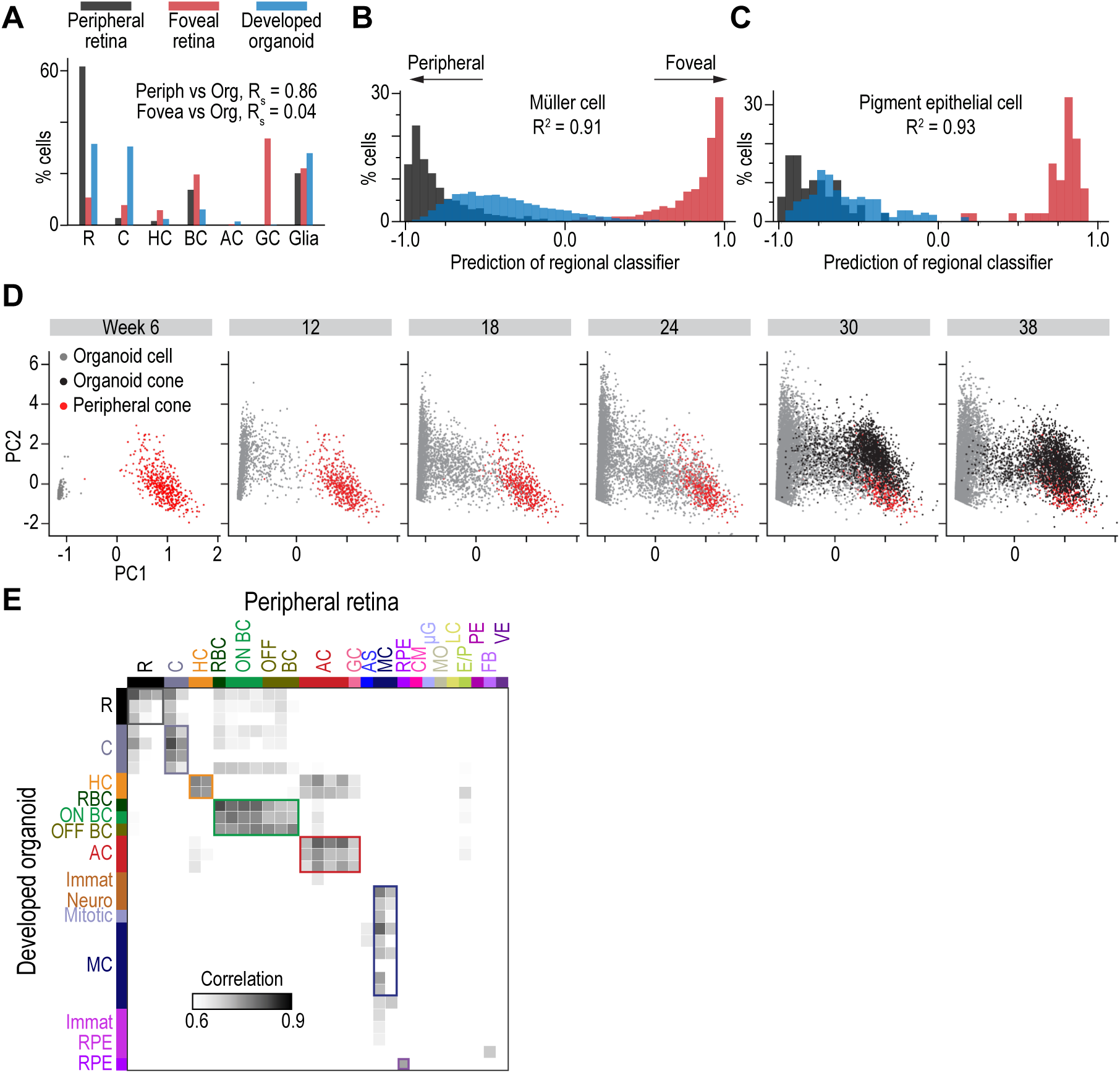
Convergence of organoid cell types to those of the adult peripheral retina. (A) Histogram of cell class composition for peripheral retina (black), foveal retina (red) and developed organoids (blue). R_S_, Spearman correlation. (B – C) Predictions from a classifier trained to predict the region of origin (peripheral, −1.0; foveal, +1.0) of (B) Müller cells and (C) pigment epithelial cells based on their transcriptome. Black bars, peripheral cells held out from training; red bars, foveal cells held out from training; blue bars, developed organoid cells; *R*^2^, coefficient of determination. (D) Comparison of transcriptomes in all organoid cells and adult cones by principal component analysis. Scatter plot axes are the first two principal components (PC1, PC2). Each point is the transcriptome of a cell. From left to right, panels contain cells from organoids of increasing age. Red, peripheral cones; black, developed organoid cones; grey, other organoid cells. (E) Correlation in gene expression between developed organoid (rows) and peripheral retina (columns); colormap at bottom left, level of correlation. Colored boxes, locations where Infomap clusters in the two tissues have the same cell class. Cell type colors and acronyms are according to Figure 5B.

We performed regression analysis to determine whether transcriptomes of adult retinal cell types differ in different regions of the retina. We developed a model (classifier) based on transcriptomes that predicted for each cell type or cell class whether single cells of a given type or class come from peripheral or foveal retina. We quantified the performance of the classifier using the coefficient of determination (R^2^), which takes values between zero and one, with higher values for more accurate classifiers and a value of one when all cells are correctly labelled as peripheral or foveal. The classifiers all had coefficients of determination above 0.25, but the highest were for Müller cells (Figure 6B; R^2^ = 0.91) and pigment epithelial cells (Figure 6C; R^2^ = 0.93) (Supplemental Figure 6; rods: R^2^ = 0.54; cones: R^2^ = 0.89; horizontal cells: R^2^ = 0.69; ON bipolar cells: R^2^ = 0.47; OFF bipolar cells: R^2^ = 0.44; amacrine cells: R^2^ = 0.46; ganglion cells: R^2^ = 0.66; astrocytes: R^2^ = 0.65; choroidal melanocytes: R^2^ = 0.56; microglia: R^2^ = 0.48; pericytes: R^2^ = 0.29; fibroblasts: R^2^ = 0.90; vascular endothelia: R^2^ = 0.52). Thus, the largest differences in gene expression between periphery and fovea are in Müller and pigment epithelial cells. The main fovea-specific genes in pigment epithelial cells included *WFDC1*, *CHI3L1*, and *CHN1* and in Müller cells *PMP2*, *CYP26A1*, and *EFEMP1*. The main periphery-specific genes in pigment epithelial cells included *TFPI2*, *FXYD3*, and *GAP43* and in Müller cells *ATP1A2*, *KCNK1*, and *MT3* (Supplemental Figure 7).

Since retinoic acid has been implicated in patterning the fovea (Silva and Cepko, 2017), we examined the cell-type specificity and regional transcript distribution of genes encoding retinoic acid catabolizing (*CYP26A1*, *CYP26C1*) or synthesizing (*ALDH1A1*, *ALDH1A3*) enzymes in adult retinas. *CYP26A1* and *CYP26C1* were expressed in foveal Müller cells (Supplemental Figure 8) and were significantly upregulated relative to the periphery (Mann-Whitney U test, *CYP26A1*: 6.8× upregulated, *P* = 6.3×10^-205^; *CYP26C1*: 52.6× upregulated, *P* = 5.3×10^-22^). *ALDH1A1* and *ALDH1A3* were present in peripheral and foveal choroidal melanocytes (Supplemental Figure 8) at a similar level (*ALDH1A1*: *P* = 0.63; *ALDH1A3*: *P* = 0.34); however, *ALDH1A3* was also expressed in pigment epithelial cells, where it was highly upregulated in the periphery relative to the fovea (817× upregulated, *P* = 4.3×10^-35^). These results confirm a higher retinoic acid sink in the fovea relative to the periphery and suggest choroidal melanocytes as a source of retinoic acid in the fovea and pigment epithelial cells as a source in the periphery.

### Convergence of cell-type transcriptomes of organoids and adult retina

Cell-type markers revealed similar cell types in developed organoids and adult retinas but did not provide a quantitative measure of how close the transcriptomes of organoid cell types were to their adult counterparts.

To address this, we first asked whether developed organoids resemble more the peripheral or the foveal retina by comparing cell-type composition across these two retinal regions. The cell-type composition of developed organoids was highly and significantly correlated with the peripheral retina, but weakly and not significantly correlated with the foveal retina (Spearman correlation (*R*_s_), periphery: *R*_s_ = 0.86, *P* = 0.01; fovea: *R*_s_ = 0.04, *P* = 0.94) (Figure 6A). When ganglion cells were excluded from the comparison, the correlation with the periphery remained but was lower (periphery: *R*_s_ = 0.83, *P* = 0.04; fovea: *R*_s_ = 0.54, *P* = 0.27), suggesting that the low number of ganglion cells in both developed organoids and peripheral retina does not fully account for the difference in the correlation.

Despite the peripheral cell-type composition of developed organoids, individual cells could still have transcriptomes with foveal characteristics. To examine this, we analyzed the regional identity of developed organoid cells using the previously described model (classifier) that successfully identified the regional origin of adult retinal cell types. The model classified the transcriptomes of developed organoid cells (Figure 6B, 6C; Supplemental Figure 6) as predominantly peripheral (> 50% peripheral, < 10% foveal, *P* < 0.05) for rods (63% with scores overlapping the peripheral cell distribution, 8% overlapping the foveal cell distribution, *P* = 1.8× 10^-307^; χ^2^ test), amacrine cells (94% peripheral, 0% foveal, *P* = 1.8×10^-307^), Müller cells (60% peripheral, 4% foveal, *P* = 1.8×10^-307^), and pigment epithelial cells (70% peripheral, 0% foveal, *P* = 4.4×10^-41^). Cones (21% peripheral, 43% foveal), ON bipolar cells (25% peripheral, 13% foveal), OFF bipolar cells (10% peripheral, 13% foveal), and horizontal cells (37% peripheral, 26% foveal) could not be classified as predominantly peripheral or foveal (≥ 10% in both regions). No cell type had a predominantly foveal regional prediction (peripheral < 10%, foveal > 50%, *P* < 0.05). Thus, in relation to cell-type composition and cell-type gene expression, developed organoids are closer to the peripheral than to the foveal retina.

We then developed two quantitative measures of closeness between groups of developed organoid cells and peripheral retinal cells. First, for a peripheral retina cell type, we identified the top 20 cell type-specific genes and performed principal component analysis (PCA) on the expression values of these genes in cells of the chosen retinal cell type as well as in cells of organoids from all developmental time points, regardless of their type. We examined the first two PCA components (PCA map) for each cell at different time points during organoid development. As an example, at week six, the characteristics of organoid cells were far away from those of cones of the adult peripheral retina (Figure 6D). With time, a group of organoid cells moved closer on the PCA map to the cloud of adult cones and by week 38 developed organoid cells intermingled with adult cones in terms of gene expression. To quantify this phenomenon, we defined organoid cells as being ‘close’ to an adult cell type if their distance from the adult distribution was below three (three SD) or five standard deviations (five SD) in the PCA map. This comparison was restricted to individual cell types such as rods, cones, Müller cells, and pigment epithelial cells. We further defined the ‘closeness’ of a particular cell type as the percentage of cells of that type in the developed organoid that were close to the adult peripheral cluster. The closeness values at week 38 were, for three SD and five SD respectively: rods, 0.4% and 0.9%; cones, 52.9% and 70.2%; Müller cells, 38.2% and 59.9%; pigment epithelial cells, 0.0% and 0.0%.

Second, we also compared pair-wise correlations between cell type averaged transcriptomes for the 1000 genes that were over-dispersed across the retina and developed organoid cell types. A strong diagonal trend was present (Figure 6E), with elevated pair-wise correlations even in rods and pigment epithelial cells that both had a low closeness. Thus, with time the spectrum of gene expression in organoid cells progressed towards that of the corresponding adult cell types and, depending on the cell type, the two distributions overlapped to different degrees. The closest overlap occurred with cones and the least overlap was recorded for pigment epithelial cells.

### Morphology of cell types in developed organoids and adult retina

We compared the morphology of cell types in developed organoids with those of the adult retina by immunostaining of standard markers for different retinal cell types and classes. Many aspects of developed organoid structure were similar to the adult retina: the outer nuclear layer contained cell bodies of rod (Figure 7A) and cone photoreceptors (Figure 7B), with cone cell bodies the most distal. Both blue cones and red/green cones were detected (Supplemental Figure 9). The inner nuclear layer of developed organoids also contained horizontal cells (Figure 7C), bipolar cells (Figure 7D), amacrine cells (Figure 7E) and Müller cells (Figure 7F). The processes of horizontal cells were elongated horizontally along the outer plexiform layer. An inner and outer limiting membrane were present (Supplemental Figure 9), with Müller cell processes contributing to their structure. Amacrine cell bodies were arranged at the proximal part of the developed organoid inner nuclear layer, matching the adult retina. Furthermore, starburst amacrine cells with processes in two strata of the inner plexiform layer (Figure 7E) were detected in developed organoids.

**Figure 7.**
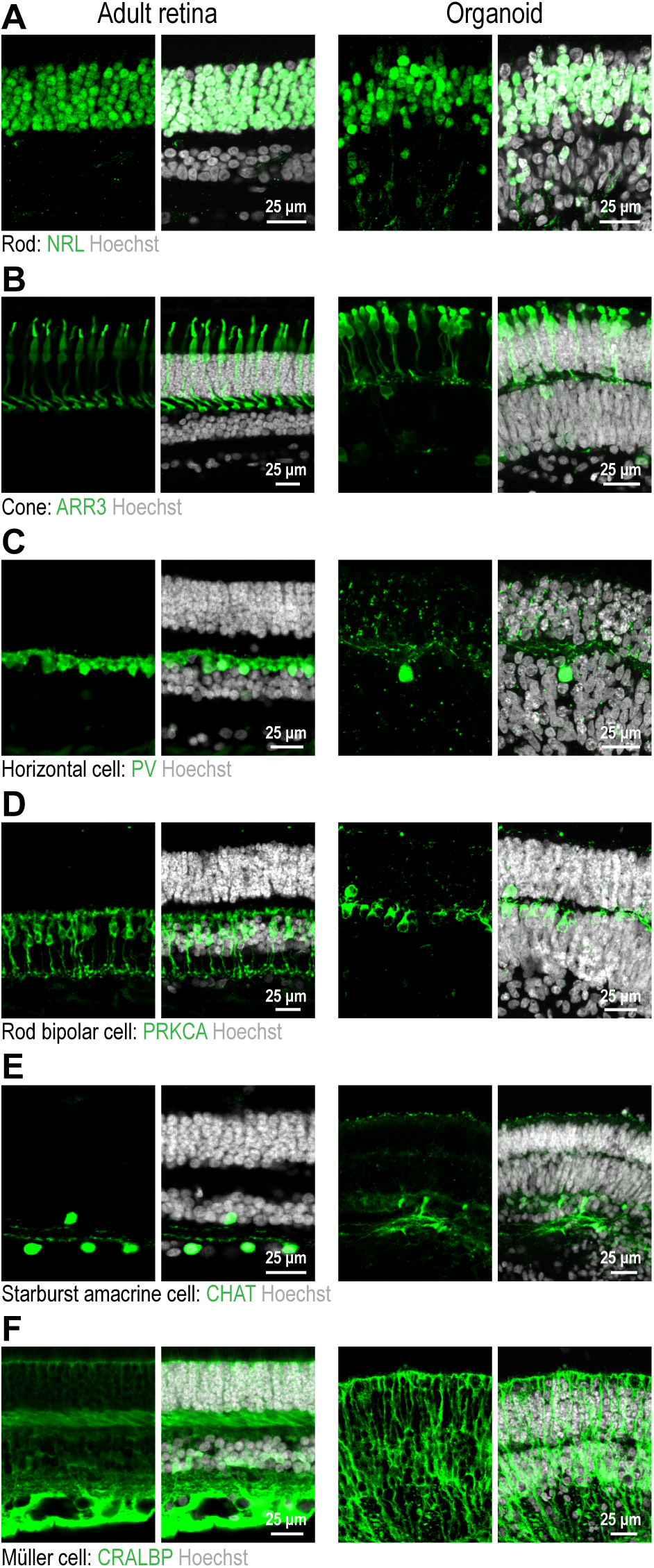
Morphology of cell types in adult human retina and organoids. (A – F) Confocal images of adult retinas (left) and organoids (right). White, Hoechst (nucleus marker). Green, antibody against (A) NRL (rod marker); (B) ARR3 (cone marker); (C) PV (horizontal cell marker); (D) PRKCA (rod bipolar cell marker); (E) CHAT (starburst amacrine cell marker); (F) RLBP1 antibody (Müller cell marker).

### Disease map of cell types in developed organoids and adult retina

We mapped genes associated with retinal disease to cell types of the adult retina and developed organoids. ‘Disease maps’ were created for 10 non-syndromic retinal diseases for the peripheral (Figure 8A; Supplemental Figure 10) and foveal retina (Supplemental Figure 10, 11) as well as for developed organoids (Figure 8B; Supplemental Figure 12): achromatopsia, congenital stationary night blindness, retinitis pigmentosa, Leber congenital amaurosis, macular degeneration, myopia, cone-rod dystrophy, choroideremia, macular dystrophy, glaucoma, and two syndromic retinal diseases Usher syndrome and Bardet–Biedl syndrome. We investigated four aspects of these disease maps.

**Figure 8.**
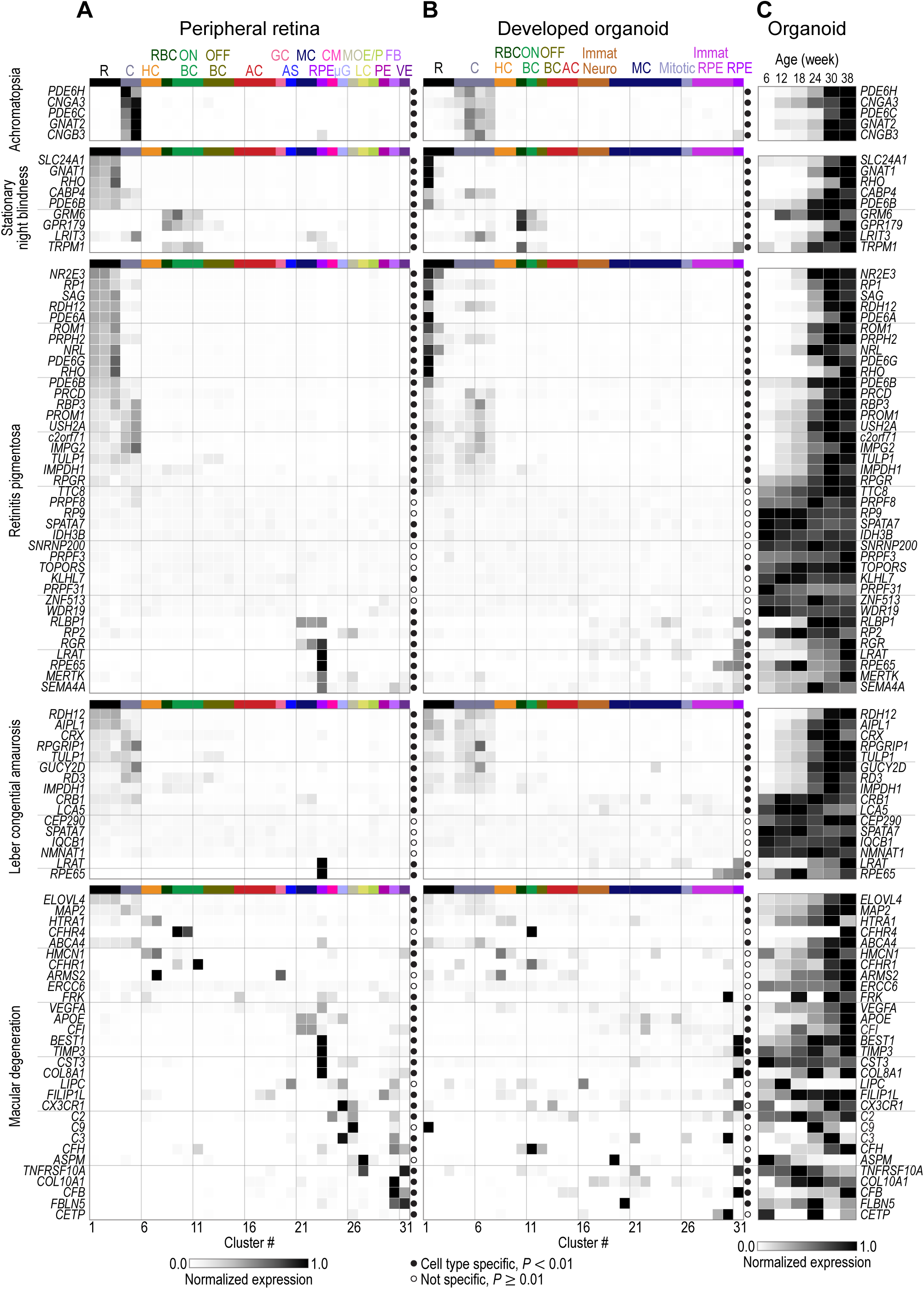
Disease map for adult human retinas and developed organoids. Normalized expression of disease genes (rows) within cell types (columns) of (A) peripheral retina and (B) developed organoid. Left, names of diseases and associated genes; colormap at bottom left of A, level of intra-gene normalized expression. Filled circle, gene significantly cell type specific (*P* < 0.01); empty circle, gene not cell type specific (*P* ≥ 0.01). Cell type colors and acronyms are according to Figure 5B. (C) Organoid age (columns) dependence of disease gene (rows) expression within organoids. Colormap at bottom, level of min-max normalized expression.

First, we asked whether diseases and disease-associated genes are cell type-specific in the adult retina. Disease specificity was defined as the uncertainty coefficient between the gene identity and cell-type origin of disease-associated transcripts, and its significance estimated by comparison to uncertainty coefficients of transcripts with shuffled cell-type labels. For individual disease-associated genes, we defined a cell-type-specificity score based on the uncertainty about the cell-type of origin for that gene’s transcripts; significance was again assessed by comparison with randomized cell-type labels.

Most of the diseases considered were cell-type specific in the peripheral and foveal retina: achromatopsia (peripheral: *P* = 5.9×10^-11^, 100% of genes significant at *P* < 0.01; foveal: *P* = 1.2×10^-10^, 100%), congenital stationary night blindness (peripheral: *P* = 7.2×10^-13^, 100%; foveal: *P* = 2.2×10^-12^, 100%), retinitis pigmentosa (peripheral: *P* = 8×10^-15^, 82%; foveal: *P* = 5.4×10^-15^, 85%), Leber congenital amaurosis (peripheral: *P* = 9.2×10^-13^, 75%; foveal: *P* = 1.1×10^-13^, 88%), macular degeneration (peripheral: *P* = 5.5×10^-13^, 80%; foveal: *P* = 1.4×10^-14^, 83%), myopia (peripheral: *P* = 1.1×10^-11^, 57%; foveal: *P* = 1.2×10^-11^, 57%), cone-rod dystrophy (peripheral: *P* = 1.7×10^-11^, 91%; foveal: *P* = 1.6×10^-11^, 88%), choroideremia (peripheral: *P* = 7.2×^10-6^, 100%; foveal: *P* = 1.2×10^-2^, 0%), macular dystrophy (peripheral: *P* = 1.5×10^-11^, 100%; foveal: *P* = 1.7×10^-11^, 100%), glaucoma (peripheral: *P* = 1.5×10^-8^, 67%; foveal: *P* = 1.4×10^-8^, 67%), Usher syndrome (peripheral: *P* = 6.7×10^-9^, 94%; foveal: *P* = 1.3×10^-7^, 100%), and Bardet-Biedl syndrome (peripheral: *P* = 1.9×10^-6^, 36%; foveal: *P* = 4.1×10^-6^, 64%;) (the significance of individual genes is indicated in Figure 8A – B; Supplemental Figure 10, 11).

Second, we asked whether the retinal cell type expressing the disease-associated gene was the same as that predicted by the clinical phenotype or expression pattern in mice. An example showing differences is the group of genes associated with age-related macular degeneration. Expression was expected and found in non-neuronal cell types such as pigment epithelial cells (*BEST1*, *P* = 6.2×10^-16^, foveal and peripheral P-values combined with Fisher’s method), microglia (*CXC3R1*, *P* = 3.0×10^-38^), and astrocytes (*APOE*, *P* = 3.4×10^-159^) (Figure 8A). Aside from photoreceptors (*ELOVL4*, *P* = 1.0×10^-32^), it was unclear whether other neuronal cell types of the human retina were involved. However, *CFH* (*P* = 5.3×10^-14^) and *CFHR1* (*P* = 3.1×10^-5^; shared locus 1q31.3) were both expressed by a type of ON cone bipolar cell negative for PCP2 (peripheral cluster #11), whereas *HTRA1* (*P* = 1.7×10^-12^) and *ARMS2* (*P* = 2.7×10^-2^; shared locus 10q26.13) were expressed in horizontal cells along with *HMCN1* (*P* = 1.0×10^-9^) (Figure 8A; Supplemental Figure 8). The 1q31.3 and 10q26.13 loci account for more disease variation than the other 32 identified macular degeneration loci combined and more than a third of the total genomic heritability (Fritsche et al., 2016). Another example was Usher syndrome. There was no previous consensus for the cell type of origin of some genes associated with type I (*USH1G*, *USH1C*, *CDH23*) and type III (*CLRN1*) Usher syndromes in the human retina as their expression varies across species (Mathur and Yang, 2015; Sahly et al., 2012; Siegert et al., 2012). We found *USH1G*, *USH1C*, *CDH23* and *CLRN1* to be highly specific for Müller cells (Supplemental Figure 10).

Third, we compared the disease maps in developed organoids and peripheral retinas (Figure 8A – B; Supplemental Figures 10, 12). The overall pattern of the disease map for most of the considered diseases was similar across developed organoids and the peripheral retina: achromatopsia (significance of specificity for developed organoids *P* = 7×10^-8^, 100% of genes significant at *P* < 0.01, 100% of genes whose cell-class with peak expression matched in developed organoids and peripheral retina), congenital stationary night blindness (*P* = 7×10^-10^, 100% specific, 100% match), retinitis pigmentosa (*P* = 3.3×10^-12^, 72% specific, 69%), Leber congenital amaurosis (*P* = 2.5×10^-10^, 75%, 75%), macular degeneration (*P* = 2×10^-12^, 63%, 45%), myopia (*P* = 8.2×10^-11^, 57%, 29%), cone-rod dystrophy (*P* = 1.7×10^-10^, 91%, 82%), choroideremia (*P* = 6.1×10^-1^, 0%, 0%), macular dystrophy (*P* = 1.5×10^-9^, 100%, 100%), glaucoma (*P* = 4.4×10^-8^, 67%, 0%), Usher syndrome (*P* = 2.2×10^-8^, 88%, 73%), and Bardet-Biedl syndrome (*P* = 1.2×10^-5^, 21%, 64%). Since the pathomechanism of the diseases associated with these genes could potentially be studied in patient-derived or gene-edited organoids, we determined the time courses of expression for these genes in organoids (Figure 8C; Supplemental Figure 12). The timing of expression was variable, with some genes constitutively expressed (e.g., *IQCB1*, *SPATA7*) but others with expression regulated up (e.g., *RHO*, *ABCA4*) or down (e.g., *ASPM*, *CLRN1*) with organoid age. Some disease-associated genes showed considerable differences in cell-type expression or degree of specificity. For example, the glaucoma-associated genes *MYOC* and *CYP1B1* are expressed in adult peripheral and foveal fibroblasts, which are absent in organoids. Instead, the two genes are expressed in developed organoid glia and pigment epithelial cells. The macular degeneration-associated gene *CX3CR1* is expressed in adult peripheral microglia but is unspecific in organoids.

Fourth, we searched for differences in the expression of disease genes across peripheral and foveal retinas that might explain the regional specificity of some retinal diseases, such as macular degeneration. For age-related macular degeneration, *CFH* and *CFHR1* (290× upregulated, *P* = 4.5×10^-80^) were expressed in bipolar cells in the periphery but not in the fovea (*CFH*, 50× upregulated, *P* = 9.1×10^-113^; *CFHR1*, 290× upregulated, *P* = 4.5×10^-80^; peripheral bipolar cell cluster #11 versus foveal cone ON bipolar cells) (Supplemental Figure 8), indicating that, in addition to their expression in pigment epithelial cells, fibroblasts and endothelia, these genes have a neuro-retinal source in the periphery. For Stargardt disease (juvenile-onset macular degeneration), the most common cause is mutations in the *ABCA4* gene (Rivera et al., 2000). Immunohistochemistry showed ABCA4 expression in photoreceptors and pigment epithelial cells of the peripheral (Figure 9A) and foveal retina (Figure 9B) as well as in photoreceptors of developed organoids (Figure 9C). It is believed that *ABCA4* expression in foveal (macular) photoreceptors underlies the specific vulnerability of the fovea. We found only a small difference in *ABCA4* expression between peripheral and foveal rods (7.2%, *P* = 0.023) and cones (19.1%, *P* = 1.7×10^-4^), with peripheral cones expressing *ABCA4* more than foveal cones (Figure 9D; Supplemental Figure 13). In contrast, the difference in *ABCA4* expression was large and highly significant between peripheral and foveal pigment epithelial cells (58%, *P* = 1.2×10^-4^) with foveal pigment epithelial cells expressing *ABCA4* more than peripheral pigment epithelial cells. Furthermore, pigment epithelial cells contained a higher percentage of *ABCA4* transcripts (0.047% of all transcripts, combining both regions) than rods (0.022%, *P* = 1.1×10^-43^) or cones (0.023%, *P* = 2.7×10^-31^) (Figure 9D; Supplemental Figure 13). Developed organoids expressed *ABCA4* primarily in rods and cones. Therefore, the macular specificity of Stargardt disease may arise from mutant *ABCA4* expression within foveal pigment epithelial cells.

**Figure 9.**
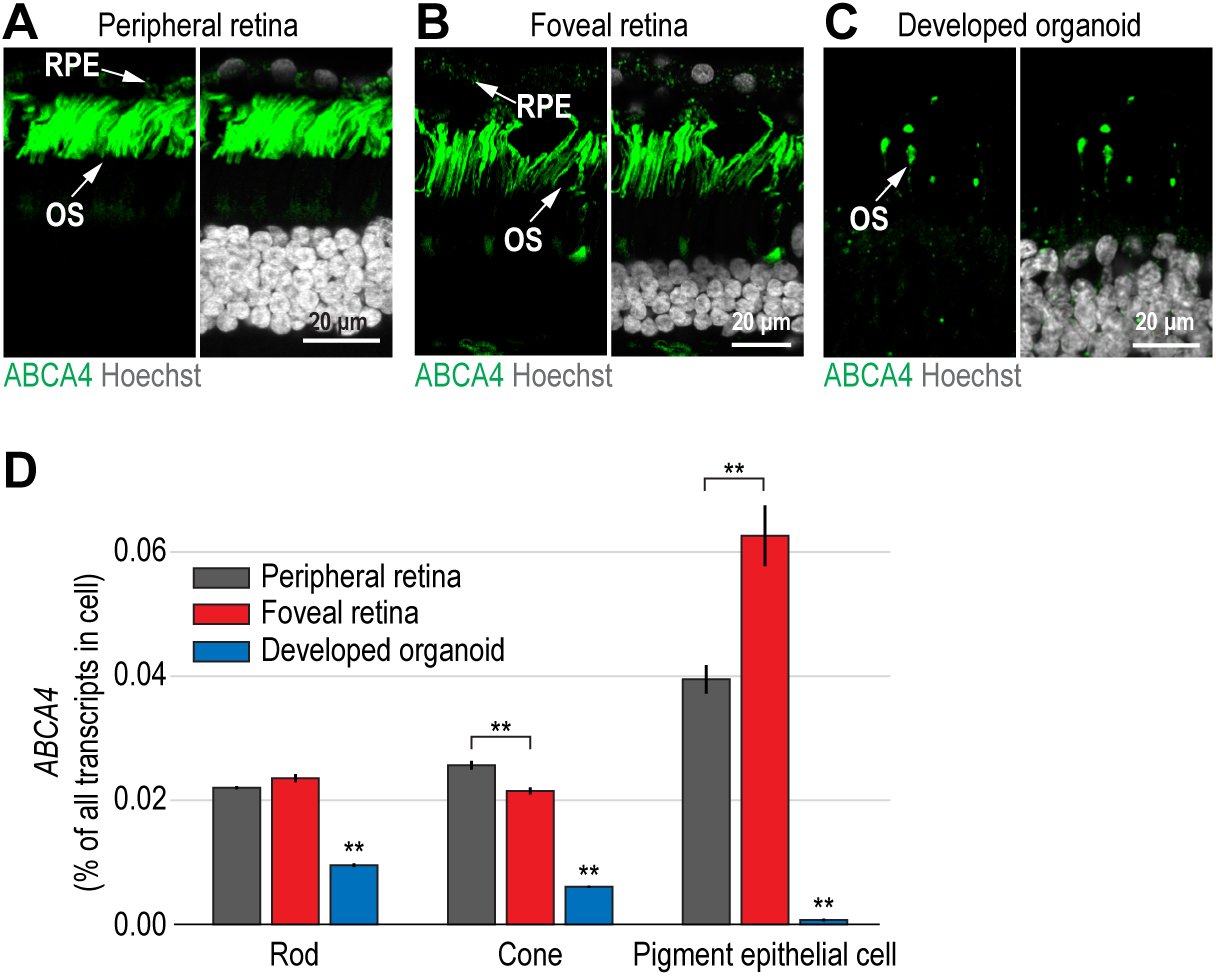
Stargardt disease gene *ABCA4* is overexpressed in foveal pigment epithelial cells but not foveal rods or cones. (A – C) Confocal images. Green, ABCA4 antibody used in (A) peripheral retina, (B) foveal retina and (C) developed organoid; white, Hoechst (nucleus marker). OS, outer segment; RPE, retinal pigment epithelial cells. (D) Expression level of *ABCA4* within cell types and regions in adult retina and developed organoids. Black, peripheral retina; red, foveal retina; blue, developed organoid. Error bars, ±1 SD. **, *P* < 1×10^-3^.

## Discussion

We have developed human retinal organoids with three nuclear and two synaptic layers and followed the development of organoid cell types using single-cell sequencing and immunohistochemistry. We found that organoid transcriptomes stabilize in a “developed” state and that the rates of organoid development *in vitro* and human retinal development *in vivo* were similar. A comparison of single-cell transcriptomes of organoids with the periphery and fovea of light-responsive adult human retinas showed that transcriptomes of organoid cell types converge to peripheral adult retina cell types. Finally, mapping disease genes to cell types of developed organoids and of adult retinas showed that inherited retinal diseases are cell-type specific and, in 67% of cases, the disease gene transcripts were detected in the same cell type or cell class in organoids and adult retinas.

### Multilayered retinal organoids

To be useful as a model system for understanding disease mechanisms and for the development of therapies, organoids should contain the relevant features and cell types of their target organ. We screened for human iPSC lines whose organoids reliably reproduced specific retinal features. We identified an iPSC line, F49B7, that generated retinal organoids with three nuclear and two synaptic layers, in which the position and morphology of retinal cell classes were similar to those in adult retina. The organoid features displayed by F49B7 suggest that screening of iPSC lines is an effective way to generate organoids with specific organ features. In addition, iPSC lines can be guided to differentiate to retinal organoids by adding growth factors (Capowski et al., 2019; Luo et al., 2018; Nakano et al., 2012). Organoids obtained with reference lines such as F49B7 will allow the study of human retina development and the effect of physical or chemical risk factors on specific cell types. For example, the impact of low oxygen or glucose on photoreceptors could be investigated in F49B7 organoids.

Given the variability in organoids produced by different iPSC lines, caution is advised when comparing organoids derived from patients with those from healthy individuals. However, gene editing within a reference cell line will allow the creation of disease models, and comparisons between organoids from mutated and reference cell lines will permit study of molecular mechanisms of diseases. For example, F49B7 could be used to study a wide variety of inherited monogenic diseases associated with photoreceptors, horizontal cells, bipolar cells, amacrine cells, Müller cells, or pigment epithelial cells.

By replacing hand microdissection of retinal structures with checkerboard scraping, we increased organoid production 160-fold. Scaling up the production of organoids with reproducible organ features and transcriptomes will allow screening of the effects of different molecules on organoid cell types. During the manual microdissection step in organoid production, many retinal structures cannot be identified by the observer and are consequently left behind. Checkerboard scraping not only removes the entire content of the plate but also avoids a key source of bias, i.e., subjective selection of retinal structures for dissection.

### Retinal organoid development at single-cell resolution

Single-cell RNA sequencing of F49B7 organoids at different time points during development revealed the successive appearance of more differentiated cells and finally all retinal cell classes. The presence of cell classes was confirmed by immunohistochemical and morphological observations. The organoid transcriptomes were reproducible (Velasco et al., 2019) at all ages from 6 to 38 weeks. By 30 weeks, organoids had stabilized and changes in gene expression between 30 and 38 weeks were less than the variability of gene expression among individual organoids of the same age. To understand how developmental age of organoids cultured *in vitro* translates to developmental age of retinas *in vivo*, we compared gene expression in developing organoids and developing human retinas at different time points. The match between organoid age and retinal age through at least week 24 was close to 1:1, which suggests that week 38 organoids closely correspond to the human retina in a newborn. The match of *in vitro* and *in vivo* human developmental rates likely generalizes to other species; for example, the fast maturation of mouse retinal organoids echoes the rapid development of mouse retina *in vivo* (Chen et al., 2016). Species-specific developmental dynamics have also been observed in cortical neurons grown from iPSCs (Otani et al., 2016).

### Functionally intact human retina

A key factor regarding the use of organoids as a model system is the similarity of cell types in organoids and cell types in the target organ. A critical factor in obtaining *in vivo*-like single-cell transcriptomes from human organs is the time cells spend in a hypoxic state after organ removal; more than 20 minutes in this state results in irreversible damage to the retina (Osborne et al., 2004). Previous single-cell RNA sequencing has been performed with human retinal cells after six hours in hypoxia (Kim et al., 2019; Peng et al., 2019). Here, we established a procedure during multi-organ donation to decrease post mortem hypoxia to less than five minutes. Our measure of health was the light-evoked spiking of retinal ganglion cells: more than 80% of recorded ganglion cells responded to light and 50% of the responding cells had a peak firing rate above 775 Hz. Critical factors in retina handling to maintain light responsivity included rapid transfer from artificial circulation *in vivo* to artificial perfusion *in vitro*, vitrectomy to improve perfusion, and reduction of lateral shear during microdissection. Notably, our recordings of light responses from 3550 ganglion cells are the first reported for a post mortem human retina and open new research avenues; for example, the use of psychophysics to link human visual perception directly to human retinal information processing (Sinha et al., 2017).

We have built a human retinal cell-type transcriptome atlas of the peripheral and foveal retina. Since the aim of our study was to provide an adult reference atlas for comparison to retinal organoids, we did not attempt to enrich the pool of dissociated cells for rare cell types (e.g., starburst amacrine cells) or classes (Peng et al., 2019). Our human retinal cell-type transcriptome atlas could be completed in the future with transcriptomes of rare cell types.

We found that the gene expression patterns of many cell types differ between the retinal periphery and the fovea, especially those of pigment epithelial cells and Müller cells. The difference in expression of two particular genes, *CYP26A1* and *ABCA4*, may be relevant for foveal biology and for the pathomechanisms of disease.

*CYP26A1*, a fovea-upregulated gene in Müller cells coding an enzyme that degrades retinoic acid has been implicated in the formation of the central, rod-free, high-acuity area in chicken (Silva and Cepko, 2017). Its presence and the foveal upregulation in human embryonic retinas (Silva and Cepko, 2017) as well as adult non-human primates (Peng et al., 2019) suggest that degradation of retinoic acid in the fovea is important, both during development and in adults. The presence of the retinoic acid-synthesizing enzymes *ALDH1A1/A3* in foveal choroidal melanocytes suggest a retinoic acid gradient between melanocytes and Müller cells and, thus, the exposure of foveal pigment epithelial cells and photoreceptor outer segments to different concentrations of retinoic acid. Understanding the role of this gradient may inform us about the microenvironment of the fovea.

The second gene, *ABCA4*, was also upregulated in the fovea. Highly significant and large (58%) foveal upregulation was only observed in pigment epithelial cells but not in rods and cones, in which *ABCA4* is also expressed. Stargardt disease, caused by mutations in the *ABCA4* gene, may thus originate in fovea-specific dysfunction of pigment epithelial cells (Lenis et al., 2018). Therapy for Stargardt disease should therefore focus on gene replacement or editing not only of photoreceptors but also of pigment epithelial cells.

### Convergence of organoid cell types to human retinal cell types

We showed that retinal organoids are transcriptomically closer to peripheral than to foveal retina. Since retinoic acid has been proposed to have a role in foveal patterning (Silva and Cepko, 2017), the presence of retinoic acid in the organoid media may contribute to their peripheral character. Although Müller cell transcriptomes are classified as peripheral in retinal organoids, they express higher levels of the fovea-specific and retinoic acid-degrading enzyme *CYP26A1* than in the fovea. This suggests active but insufficient compensation mechanisms that do not lower retinoic acid to levels conducive to fovea formation. Alternative organoid protocols exclude retinoic acid from late stages of culture and produce cone rich organoids (Kim et al., 2019). It will be interesting to evaluate whether these organoids are transcriptomically closer to the human fovea than to the periphery. Information on human fovea-specific gene expression in this resource provides insight into human foveal cell type-specific transcriptomes and can be used to validate studies attempting to induce fovea formation in retinal organoids.

Investigating whether organoid single-cell transcriptomes converge to transcriptomes of adult human cell types, we found that, at 38 weeks, about two thirds of organoid cones and Müller cells were within five SD of the type-specific gene expression of the adult cell type. Other cell types, including rods and pigment epithelial cells, were more distant but their transcriptomes were still strongly correlated with the adult cell type. Transcriptomic differences between organoid cells and adult cell types may be due to the *in vitro* culture system or simply because the developed organoids corresponded to a newborn retina and not to the adult.

### Disease map of retinal cell types

The cell-type transcriptome atlas of developed organoids and adult human retinas allows mapping of disease-associated genes to particular cell types. Single-cell transcriptome atlases for mouse (Macosko et al., 2015; Shekhar et al., 2016) and non-human primate neural retinas (Peng et al., 2019), as well as human neural retinas several hours post mortem (Kim et al., 2019; Peng et al., 2019), have been reported. The human atlas from functionally intact, five-minute post mortem neural retinas and from retinal pigment epithelium and choroid described here provides definitive information linking diseases to human cell types. The inclusion of pigment epithelium and choroid cell types is important since many retinal diseases involve these structures (Sparrow et al., 2010).

Highly significant uncertainty coefficients indicate that human retinal diseases with genetic associations are cell-type specific. Therefore, the study of a disease mechanism will be more relevant when performed in the disease-associated cell type. Moreover, the human phenotypes of some genetic diseases do not appear in model organisms such as mice (Veleri et al., 2015). Thus, the species-specific transcriptome in the relevant human cell type and genome likely matters in the pathomechanism. Since cell classes of the adult human retina are present in the developed human retinal organoid, it is potentially a model for some retinal diseases. Finally, the mapping of disease genes to cell types has implications for therapy. Therapeutic genes could be delivered to relevant cell types using adeno-associated viruses (AAVs) equipped with cell-type specific promoters (Jüttner et al., 2018).

A surprise from the disease map was the cell type of origin of gene transcripts that account for most variation in age-related macular degeneration (Fritsche et al., 2016). The *CFH*, *CFHR1* gene group at the 1q31.3 chromosomal locus had overlapping expression in a type of ON cone bipolar cells, while the *ARMS2*, *HTRA1* gene group at the 10q26.13 chromosomal locus had overlapping expression in horizontal cells. The proteins encoded by these genes are secreted. Why ON bipolar cells and horizontal cells secrete them and what role these secreted proteins play in the physiology of the retina and the pathophysiology of macular degeneration is not well understood. With the possibility of *in vivo* targeting of specific primate cell types with AAVs (Jüttner et al., 2018), and with technologies allowing individual base editing using synthetic enzymes, the primate fovea offers a platform to investigate the involvement of *CFH*, *CFHR1*, *CFHR3, ARMS2*, and *HTRA1* in fovea function and dysfunction.

Organoid cell types are not identical to human adult cell types but they do recapitulate the cell-class specificity of disease-gene expression, and the levels as well as the timing of disease-gene and gene-network expression can be examined throughout organoid development. Organoids reproduced the cell-type specificity of the diseases of the outer retina particularly well, including cone-rod dystrophy, congenital stationary night blindness, and achromatopsia. Matches were not possible for cell types absent from developed organoids (e.g., choroidal fibroblast expression of macular degeneration gene *COL10A1*).

### Human retina reference atlas

Post-mortem retinas can be obtained not only from individuals without retinal diseases but also from individuals affected by common retinal diseases such as diabetic retinopathy, age-related macular degeneration, and glaucoma. The cell-type transcriptome atlas of adult human retinas described here will serve as ground truth for future description and interpretation of the perturbations in cell-type transcriptomes obtained from patients with retinal diseases. Such comparisons could be the starting point to describe the cell-type resolved pathomechanism of diabetic retinopathy, age-related macular degeneration, and glaucoma. At least two factors are important for accurate comparison of the transcriptomes of normal and disease-affected cell types. First, since both diabetic retinopathy (Arden and Sivaprasad, 2011) and age-related macular degeneration (Blasiak et al., 2014) lead to hypoxia, the description of cell-type transcriptomes from retinas with minimal hypoxia-induced changes is critical. This is why we used retinas in this study with less than five minutes exposure to hypoxia. Second, the variation between cell-type transcriptomes from control individuals determines the number of genes for which we will be able to detect changes in gene-expression in individuals with retinal diseases. The variation across the three individuals we studied here was low for most cell types. The more post mortem retinas that can be obtained from control individuals, the more precise will be the confidence intervals described for the expression of each gene in each cell type, thus allowing more precise interpretation of changes observed in individuals with retinal diseases.

## Author contributions

M.B.S., S.O., A.H., H.P.N.S., G.R., F.N., and B.R. designed and supervised experiments. M.R., B.G.-S., M.M., J.K., C.P.P.-A., Y.H., and A.S. coordinated, optimized and performed iPSC and organoid generation. C.S.C., M.R., B.G.-S. R.C., A.W., and G.R. coordinated, optimized and performed sequencing experiments. C.S.C., D.G., N.G.-H., M.B., P.H., T.SZ., A.SZ., and A.K. coordinated, optimized, and performed retinal donations. C.S.C., R.D., and M.B optimized and performed retinal electrophysiology experiments. M.R., B.G.-S, M.M., and Y.H. optimized and performed retinal and organoid imaging. C.S.C., P.P., and S.S. performed analysis of sequencing data. C.S.C., and R.D. performed analysis of retinal electrophysiology data. C.S.C., M.R. and B.R. designed and performed statistical analyses. R.P. performed development of tools to visualize sequencing data. C.S.C., M.R., and B.R. wrote the paper.

## Acknowledgments

We thank the following people: The organ and tissue donors and their families for their generous contributions to science; Thomas Vögele, Jan Sprachta and Patricia Blaschke the transplant coordinators at the University Hospital Basel for organizing organ donations; Z. Raics and D. Hillier for developing recording software; P. Argast and P. Buchmann for electrical and mechanical engineering in support of the experiments; P. Ala-Laurila for advice on primate retinal electrophysiology. W. Baehr and A. Neutzner for sharing of antibodies. We thank FMI and NIBR core facilities for their support, especially the microscopy & imaging facility, in particular C. Genoud and A. Graff-Meyer for electron microscopy sample preparation and image acquisition. We thank the group of G. Roma for support with scRNA sequencing. We also thank P. King, F. Franke, J. Matsell, and E. Sigle for commenting on the manuscript. We thank the members of the Roska, Nigsch, Roma and Hierlemann laboratories for discussions on the manuscript. This work was supported by a Swiss National Science Foundation Synergia grant (CRSII3_141801) to A.H. and B.R.; a European Research Council Advanced Grant, a Gebert-Rüf grant, a Swiss National Science Foundation grant, the NCCR ‘Molecular Systems Engineering’ network, and a private donation from Lynn and Diana Lady Dougan to B.R.; a Swiss National Science Foundation grant, the NCCR ‘Molecular Systems Engineering’ network, a Wellcome Trust grant, a Foundation Fighting Blindness grant to H.P.N.S.

## Methods

### Reprogramming and characterization of induced pluripotent stem cells

Tissue from anonymized donors was received at 4°C in 1 ml “fibroblast medium” containing MEM alpha (Gibco, #12571), 10% fetal bovine serum (FBS; Gibco, #26140) and 100 µg / ml Primocin (Invivogen, #ant-pm-2). Biopsy pieces were rinsed with EtOH (70%) and sterile H_2_O and cultured in fibroblast medium, with media changes every other day. After 2 – 3 weeks, fibroblasts were ready to split. At 70% confluency, fibroblasts were cryopreserved in FBS (Gibco, #26140) supplemented with 20% dimethyl sulfoxide (DMSO; Fisher Scientific, #D128‒500). Fibroblasts were reprogrammed with the CytoTune-iPS Reprogramming Kit (#A13780‒01) according to the manufacturer’s instructions. In brief, 5×10^5^ fibroblasts were plated in one well of a 6-well culture plate with fibroblast medium and infected with Sendai virus at a multiplicity of infection (MOI) of 5:5:3 (KOS, MOI = 5; hc-Myc, MOI = 5; hKlf4, MOI = 3) for 16 h. On day 7, the fibroblasts were plated at 5×10^4^ – 1×10^5^ onto irradiated mouse embryonic fibroblasts (Gibco, #S1520‒100). On day 8, the medium was changed to “iPSC medium” containing Glasgow’s MEM (GMEM; Gibco, #21710‒025), 20% knockout serum replacement (Gibco, #10828.028), 1% Non-essential Amino Acid Solution (NEAA Solution; Sigma, #M7145), 2 mM GlutaMAX (Thermo Fisher Scientific, #35050038), 55 mM 2-mercaptoethanol (Gibco, #21985‒023) and 10 μg / mL of fresh Fibroblast Growth Factor-Basic (bFGF; Gibco, #PHG0264). Once iPSC colonies formed, they were picked, plated on 12-well Corning Synthemax-R plates (Corning, #3979) and transferred during the course of 4 days from iPSC medium to mTesR1 medium (STEMCELL Technologies, #85850) by mixing iPSC medium and mTesR medium (day 1, 25% mTesR; day 2, 50%; day 3, 75%; day 4, 100%).

### Culture of human induced pluripotent stem cells

iPSCs were cultured at 37°C and 5% CO_2_ in a humidified incubator in mTesR1 medium (STEMCELL Technologies, #85850) on 6-well plates (Corning, #3516) coated with Matrigel (Corning, #356230) and passaged in small clumps once per week using Versene (Gibco, #15040066) or mechanical passaging with a cell-passaging tool (Thermo Fisher Scientific, #23181010) according to WiCell protocols (https://www.wicell.org/stem-cell-protocols.cmsx). Cells tested negative for mycoplasma on a regular basis using the MycoAlert PLUS Mycoplasma Detection Kit (Lonza, #CABRLT07‒710). Cell identity was confirmed by short tandem repeat analysis (STR analysis; Microsynth, Switzerland) on genomic DNA extracted from fibroblasts and iPSCs by the DNeasy Blood & Tissue Kit (Qiagen, #69504).

### Characterization of human induced pluripotent stem cells

iPSC pluripotency was assessed by staining iPSC colonies cultured in 6-well plates for alkaline phosphatase using the Vector Blue Alkaline Phosphatase Kit (Reactorlab, #SK‒5300), 4 – 6 days after splitting. iPSCs seeded into Matrigel-coated (Corning, #356230) 8-well chamber slides (ibidi, #80826) were further stained for the pluripotency markers SOX2, OCT4, NANOG and SSEA4 (additional antibody information is in the table “List of antibodies used for immunostainings”). The potential to differentiate into the three embryonic germ layers was assessed by directed differentiation into ectoderm, mesoderm and endoderm using the STEMdiff Trilineage Differentiation Kit (STEMCELL Technologies, #05230) according to the manufacturer’s instructions on cells maintained in Matrigel-coated (Corning, #356230) 8-well chamber slides (ibidi, #80826). To test for chromosomal aberrations, G-banded karyotyping was performed on 20 cells per line (Cell Guidance Systems, UK). Further, array comparative genomic hybridization (array CGH) was performed using the Illumina HumanOmni2.5Exome-8 BeadChip v1.3 (LIFE & BRAIN GmbH, Germany) on genomic DNA extracted from iPSCs by the DNeasy Blood & Tissue Kit (Qiagen, #69504).

### Generation of retinal organoids from human induced pluripotent stem cells

Retinal organoids were generated as described before (Zhong et al., 2014) with some modifications. On day zero of differentiation, floating embryoid bodies (EBs) were generated by dissociating iPSC colonies from one well of a 6-well plate (Corning, #3516) into evenly sized clumps with a cell-passaging tool (Thermo Fisher Scientific, #23181010). The EBs were cultured in suspension in mTesR1 medium supplemented with 10 µM blebbistatin (Sigma, #B0560‒5MG) on 3.5-cm untreated petri dishes (Corning, #351008). On days 1 and 2, one third of the medium was exchanged with “neural induction medium” (NIM) containing DMEM/F12 (Gibco, #31331‒028), 1× N2 Supplement (Gibco, #17502‒048), 1% NEAA Solution (Sigma, #M7145) and 2 µg / ml heparin (Sigma, #H3149‒50KU). On day 3, aggregates were sedimented by gravity in a 15-ml tube, washed with NIM, and cultured in a 3.5-cm untreated petri dish (Corning, #351008) in NIM. Half of the NIM was exchanged daily. On day 7, aggregates from one 3.5-cm dish were plated onto a 6-cm dish (Corning, #430166) coated with Growth Factor-Reduced Matrigel (Corning, #356230) and then maintained with daily NIM changes. On day 16, NIM was exchanged for “3:1 medium” containing 3 parts DMEM (Gibco, #10569‒010) per 1 part F12 medium (Gibco, #31765‒ 027), supplemented with 2% B27 without vitamin A (Gibco, #12587-010), 1% NEAA Solution, 1% penicillin/streptomycin (Gibco, #15140‒122). On day 28 – 32 retinal structures were detached from the Matrigel plate by checkerboard scraping. First, to break the tissue sheets into smaller pieces, a 1 to 2-mm^2^ grid was scratched through the cells on the culture plate with a 10-µl or 200-µl pipette tip. Second, the entire contents of the culture plate were washed off the plate with a 1000-µl pipette tip to generate numerous retinal aggregates and small uncharacterized debris. Retinal structures were typically not broken by the pipette passing through the dish, but rather came off the plate as large tissue pieces. The aggregates contained both regions of neural retina and retinal pigment epithelium. To remove debris and single cells, the aggregates were washed 3× in a 15-ml tube by sedimentation in 3:1 medium and then maintained in suspension on sterile petri dishes (VWR, #391‒2016) in 10 – 15 ml 3:1 medium with media changes every other day. Aggregates without phase-bright, stratified neuroepithelium (Supplemental Figure 2) were sorted out one week after scraping to leave behind only high-quality retinal organoids. Even before sorting, up to 80% of aggregates within the plate contained phase-bright, stratified neuroepithelium.

From day 42, aggregates were cultured in 3:1 medium supplemented with an additional 10% heat-inactivated FBS (Millipore, #es‒009‒b) and 100 µM taurine (Sigma, #T0625‒25G) with media changes every other day. At week 10, the culture medium was supplemented with 10 µM retinoic acid (Sigma, #R2625). From week 14, the B27 supplement in 3:1 media was replaced by N2 supplement (Gibco, #17502‒048) and retinoic acid was reduced to 5 µM.

### Primary human tissue samples

Human retina tissue was obtained from multi-organ donors by sampling non-transplantable eye tissue that was removed in the course of cornea harvesting for transplantation. Donors with a documented history of eye disease were excluded from this study. Personal identifiers were removed and samples were coded before processing. All tissue samples were obtained in accordance with the tenets of the Declaration of Helsinki, and all experimental protocols were approved by the local ethics committee. The sequencing results include *n* = 6 eyes from *n* = 3 donors. One donor was male and two were female.

### Human retinal tissue handling

Eye enucleation was performed while circulation and ventilation were still in place. The enucleated eye was opened and the eye cup flattened with butterfly cuts. The vitreous was then removed. The tissue was submerged in flowing Ames medium (Sigma, #A1420), saturated with carbogen gas (95% O_2_, 5% CO_2_). The time elapsed from central retinal artery clamp to artificial *ex vivo* perfusion was less than 5 minutes for all eyes. Samples intended for electrophysiology were subsequently handled in a dark room using dim red illumination for general dissection and night vision (ATN Corp, USA) with infrared illumination for dissection under a microscope. The regions of the retina sampled for sequencing were: (i) peripheral retina; square 2 – 3 mm on a side; ventral midline at 50% eccentricity (midway between the fovea and ventral retina border with the ora serrata) (ii) foveal retina; disc 1.5 mm in diameter; centered on the fovea centralis. Landmarks used to locate the fovea included (i) adjacency to the optic disc, (ii) the avascular zone, (iii) pigmentation of the macula lutea, and (iv) retinal transparency at the fovea centralis due to retinal thinning. In both regions, the neural retina tissue was separated from the retinal pigment epithelium/choroid and the two tissues were dissociated separately.

### Immunofluorescence staining and imaging

Human retina tissue was fixed over night at 4°C in 4% weight / volume (w / v) paraformaldehyde (PFA) in phosphate buffered saline (PBS). Organoids were fixed for 4 h at 4°C in 4% PFA in PBS. After fixation, samples were washed 3× 30 min with PBS and cryopreserved in 30% sucrose in PBS over night at 4°C. Samples were stored at −80°C until use.

Cryosections (20 – 40 μm) were generated using a cryostat (MICROM International, #HM560) on organoids and human retina embedded in OCT compound (VWR, #25608‒930). Sections were mounted onto Superfrost Plus slides (Thermo Fisher Scientific, #10149870), dried for 4 to 16 h at room temperature and stored at −80°C until use. Photoreceptor outer segments in retinal organoids were not preserved upon OCT embedding. Therefore, for cryosectioning of photoreceptors, organoids were embedded in 7.5% gelatin and 10% sucrose in PBS (Lancaster and Knoblich, 2014b).

For immunostainings of cryosections, slides were first dried for 30 minutes at room temperature and then rehydrated for 5 – 10 min in PBS. Second, slides were blocked with “blocking buffer A” which was PBS supplemented with 10% normal donkey serum (Sigma, #S30‒100ML), 1% (w / v) bovine serum albumin (BSA; Sigma, #05482‒25G), 0.5% Triton X-100 in PBS (Sigma, #T9284‒500ML) and 0.02% sodium azide (Sigma, #S2002‒25G) at room temperature for 1 h. Sections were then incubated in a humidified chamber with primary antibodies (see table, “List of antibodies used for immunostainings”). For each slide, primary antibodies were diluted in 100 µl of “blocking buffer B” which was PBS supplemented with 3% normal donkey serum, 1% BSA, 0.5% Triton X-100 in PBS and 0.02% sodium azide over night at room temperature. After washing 3× 15 min in PBS with 0.1% TWEEN 20 (Sigma, #P9416‒100ML), slides were incubated with secondary antibodies (Thermo Fisher Scientific, donkey secondary antibodies conjugated to Alexa Fluor 488, 568 or 647) diluted 1:500 in 100 µl blocking buffer B at room temperature in the dark. The sections underwent washes of 2× 15 min in PBS with 0.1% Tween, one wash of 15 min in PBS, and were cover-slipped with ProLong Gold (Thermo Fisher Scientific, #P36934). Images were acquired with an LSM 700 confocal microscope (Zeiss) or a spinning disc microscope (Axio Imager M2 upright microscope, Yokogawa CSU W1 dual camera T2 spinning disk confocal scanning unit, Visitron VS-Homogenizer).

For Vibratome sectioning (VT1000S vibratome, Leica), organoids were embedded in 3% agarose (Promega, #V3125), sectioned at 120-µm thickness and collected in 30% sucrose in 1× PBS for cryopreservation. Sections were stored at −80°C until use Vibratome sections were stained floating in 24-well plates by incubating for 5 h in blocking buffer A at room temperature followed by incubation in 120 µl primary antibody solution diluted in blocking buffer B for 3 to 6 days at 4°C. For vibratome sections, the antibody concentrations were doubled compared to those described for cryosections. Sections were washed 3× in PBS supplemented with 0.05% Triton X-100 (PBS-T) for a total of 24 h, incubated with secondary antibodies (Thermo Fisher Scientific, donkey secondary antibodies conjugated to Alexa Fluor 488, 568 or 647), diluted 1:250 in blocking buffer B for 4 h at room temperature and washed 3× in PBS-T for a total of 24 h. Sections were transferred to glass slides using a paint brush and mounted using ProLong Gold (Thermo Fisher Scientific, #P36934). Images were acquired using a spinning disc microscope (Nikon Ti2-E Eclipse inverted, Yokogawa CSU W1 dual camera T2 spinning disk confocal scanning unit, Visitron VS-Homogenizer).

Wholemount staining of organoids were performed by incubating fixed organoids for 3 days in blocking buffer A at room temperature, followed by incubation in 100 µl primary antibody solution diluted in blocking buffer B for 5 days at 4°C. For wholemount staining, the antibody concentrations were double those described for cryosections. Organoids were washed 3× in PBS supplemented with 0.05% Triton X-100 for a total of 3 days at room temperature, incubated with secondary antibodies (Thermo Fisher Scientific, donkey secondary antibodies conjugated to Alexa Fluor 488, 568 or 647), diluted 1:250 in blocking buffer B for 4 days at 4°C and washed 3× in PBS-T for a total of 4 days. Organoids were imaged on an inverted spinning disk confocal microscope (Nikon Ti2-E Eclipse Inverted motorized stand + Yokogawa CSU W1 Dual camera T2 spinning disk confocal scanning unit. Visitron VS-Homogenizer) in 8-well chamber slides (ibidi, #80826).

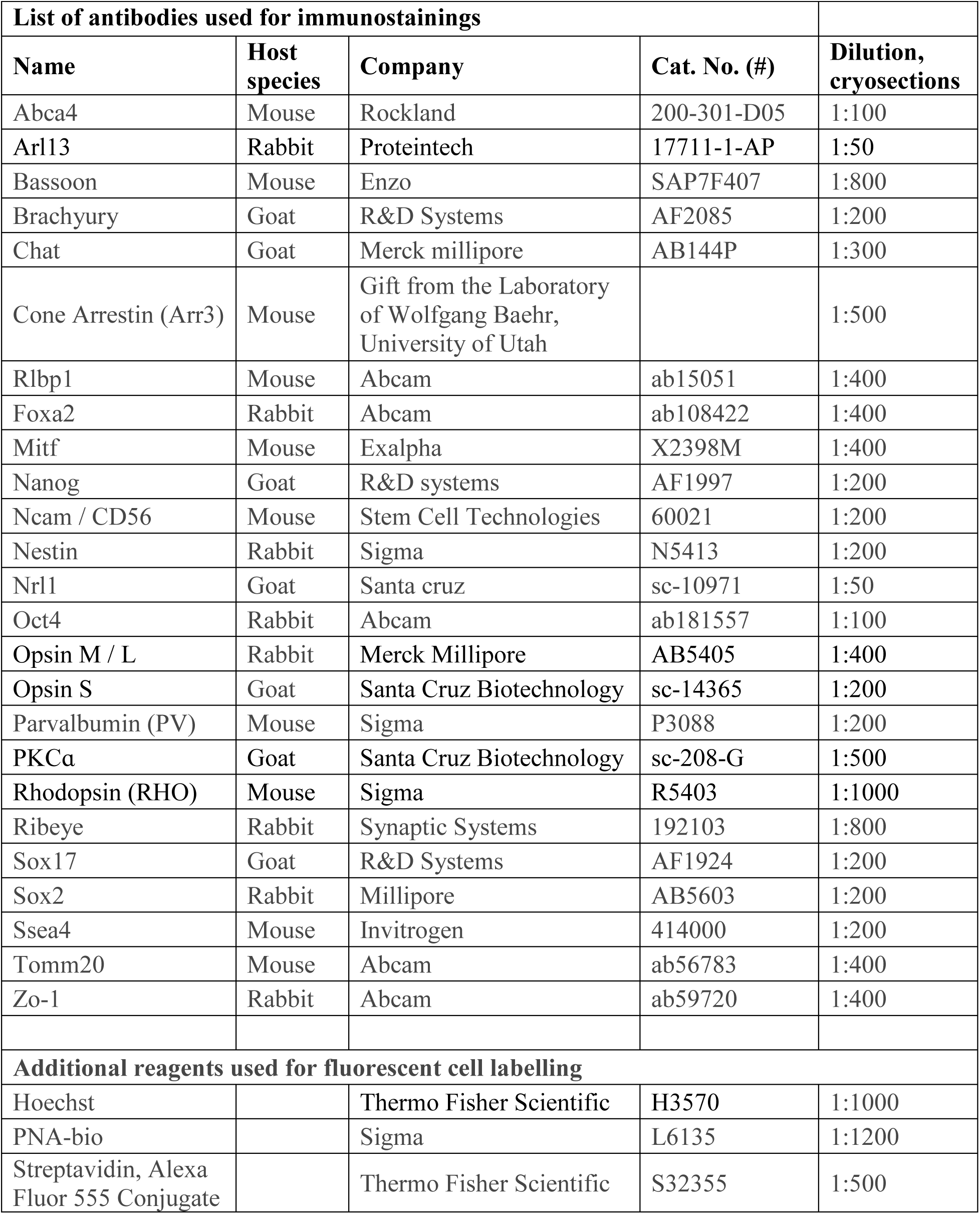

| **List of antibodies used for immunostainings** |  |  |  |  |
| --- | --- | --- | --- | --- |
| **Name** | **Host species** | **Company** | **Cat. No. (#)** | **Dilution, cryosections** |
| Abca4 | Mouse | Rockland | 200-301-D05 | 1:100 |
| Arl13 | Rabbit | Proteintech | 17711-1-AP | 1:50 |
| Bassoon | Mouse | Enzo | SAP7F407 | 1:800 |
| Brachyury | Goat | R&D Systems | AF2085 | 1:200 |
| Chat | Goat | Merck millipore | AB144P | 1:300 |
| Cone Arrestin (Arr3) | Mouse | Gift from the Laboratory of Wolfgang Baehr, University of Utah |  | 1:500 |
| Rlbp1 | Mouse | Abcam | ab15051 | 1:400 |
| Foxa2 | Rabbit | Abcam | ab108422 | 1:400 |
| Mitf | Mouse | Exalpha | X2398M | 1:400 |
| Nanog | Goat | R&D systems | AF1997 | 1:200 |
| Ncam / CD56 | Mouse | Stem Cell Technologies | 60021 | 1:200 |
| Nestin | Rabbit | Sigma | N5413 | 1:200 |
| Nrl1 | Goat | Santa cruz | sc-10971 | 1:50 |
| Oct4 | Rabbit | Abcam | ab181557 | 1:100 |
| Opsin M / L | Rabbit | Merck Millipore | AB5405 | 1:400 |
| Opsin S | Goat | Santa Cruz Biotechnology | sc-14365 | 1:200 |
| Parvalbumin (PV) | Mouse | Sigma | P3088 | 1:200 |
| PKCa | Goat | Santa Cruz Biotechnology | sc-208-G | 1:500 |
| Rhodopsin (RHO) | Mouse | Sigma | R5403 | 1:1000 |
| Ribeye | Rabbit | Synaptic Systems | 192103 | 1:800 |
| Sox17 | Goat | R&D Systems | AF1924 | 1:200 |
| Sox2 | Rabbit | Millipore | AB5603 | 1:200 |
| Ssea4 | Mouse | Invitrogen | 414000 | 1:200 |
| Tomm20 | Mouse | Abcam | ab56783 | 1:400 |
| Zo-1 | Rabbit | Abcam | ab59720 | 1:400 |
| **Additional reagents used for fluorescent cell labelling** |  |  |  |  |
| Hoechst |  | Thermo Fisher Scientific | H3570 | 1:1000 |
| PNA-bio |  | Sigma | L6135 | 1:1200 |
| Streptavidin, Alexa Fluor 555 Conjugate |  | Thermo Fisher Scientific | S32355 | 1:500 |

### Electron microscopy

Organoids were fixed in 2.5% glutaraldehyde (Electron Microscopy Sciences, #16220) and 2% PFA (Electron Microscopy Sciences, #15710) in 0.1 M sodium cacodylate buffer (Sigma, #20840) overnight at 4°C, washed with 0.1 M cacodylate buffer, and post-fixed in 1% osmium tetroxide (Electron Microscopy Sciences, #19170) for 2 h at room temperature. Organoids were then dehydrated in a graded ethanol (Sigma, #51976) series (25%, 50%, 75%, and 100%), further dehydrated in propylene oxide (Sigma, #82320) and embedded in Epon resin (SERVA Electrophoresis; glycid ether #21045.01, dodecenylsuccinic acid anhydride #20755.01, methylnadic anhydride #29452.01 and benzyl dimethylamine #14835.01) for 12 h. Semi-thin (0.4 µm) and ultra-thin sections (50 nm) were cut with a Leica EM UC7 ultramicrotome and the latter were collected on formvar-coated single-slot copper grids (Electron Microscopy Sciences, #FF2010-Cu) for imaging. Sections were contrasted with 1% uranyl acetate (Electron Microscopy Sciences, #22400) and lead citrate (Electron Microscopy Sciences, #17800) (8 min each) and imaged on a FEI Tecnai Spirit electron microscope (FEI Company) operated at 120 kV using a side-mounted 2K × 2K CCD camera (Veleta, Olympus).

### Dissociation of organoids and human retina into single cells

The retinal region of organoids, which could be clearly distinguished in light microscopy, was dissected from the main body of the organoid using needles. The single-cell dissociation protocol was adapted from an existing protocol (Balse et al., 2005). The retinal piece (derived from human retina or retinal organoid) was washed 2× at 37°C with 1 ml “ringer solution” without calcium containing 125.6 mM NaCl, 3.6 mM KCl, 1.2 mM MgCl_2_·H_2_O, 22.6 mM NaHCO_3_, 21.7 μM NaH_2_PO_4_·H_2_O, 70.2 μM NaH_2_PO_4_·2 H_2_O, 1.2 mM Na_2_SO_4_, 10.0 mM D-Glucose and 0.4 mM EDTA. “Activated papain solution” was prepared by mixing 8 U of papain (Worthington Biochemical, #LS003127) with 48 µl of “activator” containing H_2_0 with 1.1 µM EDTA, 5.5 mM L-cysteine and 0.07 mM 2-mercaptoethanol and incubating for 30 min at 37°C before diluting with 950 µl of 37°C ringer solution. Tissue pieces were incubated at 37°C in activated papain solution: 300 µl per organoid for 35 min; 500 µl per human retinal piece for 30 min. The papain digestion reaction was stopped by placing the tubes on ice and adding equal volumes of “stop solution” containing Neurobasal A medium (Thermo Fisher Scientific, #10888022), 2 mM GlutaMAX (Thermo Fisher Scientific, #35050038), 10% FBS (Millipore, #es‒009‒b) and 20 U / ml DNAse (Sigma, #D4263). Samples were centrifuged at 200 *g* and 4°C for 30 s and washed using Neurobasal A supplemented with 2 mM GlutaMAX and 10% FBS: 1 ml per organoid; 1.5 ml per retinal piece. A single cell suspension was generated by gently triturating 20× with a 1000-µl pipette tip in ice cold Neurobasal A (Thermo Fisher Scientific, #10888022), 2 mM GlutaMAX (Thermo Fisher Scientific, #35050038), 2% B27 supplement without vitamin A (Gibco, #12587‒010), and 20 U / ml DNAse (Sigma, #D4263) or until no large clumps were visible: 300 µl per organoid; 500 µl per human retinal piece. The cells were collected by centrifuging 5 min at 300 *g* and 4°C and resuspended in PBS containing 0.04% BSA (Thermo Fisher Scientific, #AM2616). The cell suspension was filtered to remove cell aggregates and particles with a diameter greater than 35 µm. Cells were mixed 1:1 with trypan blue (Thermo Fisher Scientific, #T10282) and counted using the Countess II cell counter (Thermo Fisher Scientific); cell viability was typically above 80%.

### Single cell RNA-Sequencing

Cellular suspensions (8000 cells per lane) were loaded on a 10x Genomics Chromium Single Cell instrument to generate single-cell Gel Beads in Emulsion (GEMs). Single-cell RNA-Seq libraries were prepared using GemCode Single Cell 3’ Gel Bead and Library Kit according to the manufacturer’s manual (version CG00052_SingleCell3’ReagentKitv2UserGuide_RevD). Reverse transcription of GEMs was performed in a Bio-Rad PTC-200 thermal cycler with a semi-skirted 96-well plate (Eppendorf, #0030 128.605): 53°C for 45 min, 85°C for 5 min, held at 4°C. After reverse transcription, the GEMs emulsion was broken and the single-stranded cDNA was cleaned up with DynaBeads MyOne Silane Beads (Life Technologies, #37002D). cDNA was amplified using a Bio-Rad PTC-200 thermal cycler with 0.2-ml 8-strip non-flex PCR tubes with flat caps (STARLAB, #I1402-3700): 98°C for 3 min, cycled 12×: 98°C for 15 s, 67°C for 20 s, and 72°C for 1 min; 72°C for 1 min, held at 4°C. Amplified cDNA product was cleaned up with the SPRIselect Reagent Kit (0.6× SPRI; Beckman Coulter, #B23317). Indexed sequencing libraries were constructed using the reagents in the Chromium Single Cell 3’ library kit V2 (10x Genomics, #120237), following these steps: (i) fragmentation, end repair and A-tailing, (ii) post fragmentation, end repair and A-tailing; double-sided size selection with SPRIselect Reagent Kit (0.6× SPRI and 0.8× SPRI), (iii) adaptor ligation, (iv) post-ligation cleanups with SPRIselect (0.8× SPRI), (v) sample index PCR using the Chromium multiplex kit (10x Genomics, #120262), and (vi) post-sample index double-sided size selection with SPRIselect Reagent Kit (0.6× SPRI and 0.8× SPRI). The barcode sequencing libraries were measured using a Qubit 2.0 with a Qubit dsDNA HS assay kit (Invitrogen, #Q32854) and the quality of the libraries assessed on a 2100 Bioanalyzer (Agilent) using a high-sensitivity DNA kit (Agilent, #5067‒4626). Sequencing libraries were loaded at 10 – 12 pM on an Illumina HiSeq2500 with 2 × 50 paired-end kits using the following read length: 26 cycles Read1, 8 cycles i7 Index, and 98 cycles Read2.

### Microelectrode array recording and visual stimulation of human retinas

The neural retina and pigment-epithelium/choroid, which had strong interconnections, were left attached to each other until a sample was recorded. In this manner, light responses could be recorded from samples of artificially perfused *ex vivo* human retina for up to 16 h post mortem. Before recording, a small piece of retina (3 mm × 3 mm) was cut and the neural retina isolated from the pigment epithelium and choroid. The neural retina was placed onto a high-density microelectrode array and stabilized using a transparent permeabilized membrane (Polyester, 10-µm thickness, 200-µm hole size, 400-µm spacing). The high-density microelectrode array had the dimensions 3.85 × 2.1 mm and contained 26,400 platinum electrodes, of which 1,024 could have their electrical activity recorded simultaneously (Müller et al., 2015). A chamber surrounding the electrode array was perfused with Ames media (Sigma, #A1420) saturated with carbogen gas at 37°C. During recording, patterned light stimulation was delivered. Stimuli were generated by a DLP projector (K10, Acer) with optics modified to project onto the retinal surface. The projector was controlled by custom Python software developed by Z. Raics. The stimulus was a light step followed by a frequency chirp and an intensity sweep (Baden et al., 2016). Maximal light irradiance was 29.3 µW / mm^2^ in the plane of the retina and white light was generated using achromatic RGB triplets.

### Quantification and statistical analysis

#### Electrophysiology analysis

The raw electrical activity on the array was spike sorted using an automatic spike sorter dedicated to high-density microelectrode arrays to identify sets of action potentials arising from the same cell. Because individual spikes were detected on multiple electrodes, the sorter could cluster them using the combined information of the shapes and spatial extent of their waveforms (Diggelmann et al., 2018). Retinas were stimulated with a temporally repeating pattern of spatially uniform white light. Each repetition of the stimulus was considered a trial *Q*_*k*_ and an entire experiment *Q* consisted of *n*_trial_ trials. Spikes from each 30-s trial were binned into *n*_bin_ = 120 bins of width 250 ms to create the matrix **ρ** with dimensions *n*_bin_ × *n*_trial_ whose elements contain the spike counts of each bin in each trial.

Light responsivity was assessed by correlating spike counts between all possible combinations of pairs of trials at once. For example, when *n*_trial_ = 3, the number of possible combinations of pairs of trials is 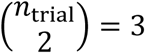. These three pairs are 〈(1,2), (1,3), (2,3)〉. The vectors containing the binned spike counts for these three pairs of trials would then be

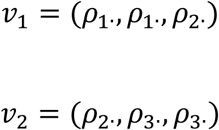

with *ρ*_*i*·_ = (*ρ*_*i*1_, *ρ*_*i*2_, …, *ρ*_*in*_bin__) the binned spike counts for each trial *i*.

Cells were evaluated if their average firing rate exceeded 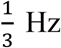 (*n* = 4073 of 4343 cells) and considered light responsive if their Pearson correlation coefficient *R*_P_(*v*_1_, *v*_2_) was positive and significant at *P* < 0.01 (*n* = 3350 of 4073 cells). The reported peak firing rate of a cell was the inverse of the average of the 5 shortest inter-spike intervals across the experiment.

Light responses were clustered by the Infomap method. First, *K*-nearest neighbors were approximated (Annoy v1.15.2, Spotify) using 200 trees and *K* = 60. Second, the graph was clustered by Infomap using the python-igraph library. For visualization, light responses were embedded using an adaptation of the scVis algorithm (Ding et al., 2018) for electrophysiology data. Briefly, the t-SNE objective function was implemented inside a variational autoencoder (InfoVAE) (Zhao et al., 2017) implemented in Keras, using the Tensorflow backend, and trained with a perplexity of 30.

#### Single-cell RNA sequencing analysis

##### Read alignment and expression count table generation

The CellRanger package (v2.0, 10x Genomics) was used to extract unique molecular identifiers, cell barcodes, and genomic reads from the sequencing results of 10x Chromium experiments. Reads were aligned against the annotated human genome (GRCh38, GENCODE v27), including both protein coding and non-coding transcripts. Reads with the same cell barcode and unique molecular identifier were collapsed to a unique transcript. In order to discard potentially unhealthy or damaged cells, the built-in cell filtering step of CellRanger was followed by further cell filtering using as criteria the number of detected transcripts and the fraction of mitochondrial reads. For both criteria, a LOESS fit was performed on the corresponding feature’s rank-size distribution; the slope of the fitted curve was determined at steps of relative width 1 / 10,000. The threshold was then set at inflection points past the 5^th^ (number of detected genes) or 95^th^ (fraction of mitochondrial reads) percentiles of the distributions. On average, an additional 2.5% of the cells were removed at this step. Transcripts from mitochondrial- and ribosomal-protein coding genes, which are typically very highly expressed and highly dispersed, irrespective of biological identity, were also discarded prior to subsequent analyses.

Two random variables were defined on the sample space of unique transcripts: the ‘cellular identity’ and ‘genetic identity’. Letting *κ* be the total number of unique transcripts, the random variable cellular identity (*C*) groups transcripts with matching cell barcodes into mutually disjoint subsets *C*_1_, *C*_2_, …, *C*_*n*_cell__ where *n*_cell_ is the number of single cells and the cardinality |*C*_*i*_| is the number of transcripts in cell *C*_i_ such that 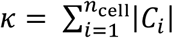. Formally, the values of *C* are 1, 2, …, *n*_cell_ corresponding to the subsets *C*_1_, *C*_2_, …, *C*_*n*_cell__. Similarly, the random variable genetic identity (*G*) groups transcripts together that belong to the same gene. The values of *G* are the mutually disjoint subsets *G*_1_, *G*_2_, …, *G*_*n*_gene__ where *n*_gene_ is the total number of genes in the annotation. *G*_*j*_, therefore, represents a particular gene and contains each transcript of this gene. If the cardinality |*G*_*j*_| is the total number of transcripts in *G_j_* then 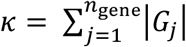. The contingency table between *C* and *G* is the count matrix **M** whose elements *M*_*ij*_ are the number of transcripts observed in cell *C*_*i*_ and gene *G*_*j*_.

##### Embedding transcriptomes into maps with scVis

Let *μ*_*G_j_*_ and *σ*_*G_j_*_ denote the mean and standard deviation of the transcript counts for gene *G*_*j*_ across all cells and *α* = 0.8 be a trend adjustment that was empirically estimated on held-out data. The trend-adjusted log coefficient of variation was

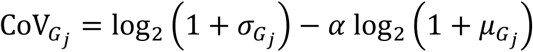

Adding 1 before taking the logarithm preserved matrix sparsity, kept values positive, and flattened the gradient for the smallest expression values. The 1000 genes with the highest CoV_*G_j_*_ were retained for dimensionality reduction. Gene selection was performed separately for organoids, developed organoids and adult retina.

The average number of transcripts per cell in the sample was

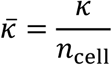

Let *M*_*i*·_ = (*M*_*i*1_, *M*_*i*2_, …, *M*_*in*_gene__) denote the vector of per-gene transcript counts for cell *C*_*i*_ and recall that |*C*_*i*_| is the total transcript count for cell *C*_*i*_. We define the scaled normalized expression vector as

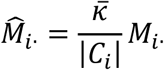

Because the expression values were non-negative 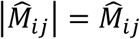 and the normalized expression’s *L*_1_ norm 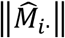 is equal to its sum.

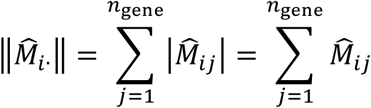

After normalizing, the sum of the expression in each cell was the same 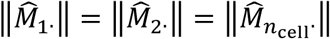 and equal to 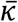, the average transcript count per cell before normalizing. The normalized expression values were then converted to a logarithmic scale, 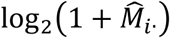, and incremental principal component analysis was performed to estimate scores for the top 100 principal components.

Before the final dimensionality reduction step, cells with very few transcripts were removed, in response to observations that their additional noise negatively influenced the learned embedding. Cells were split into a high- and a low-expression pool based on |*C*_*i*_|. Thresholds for high expression were – peripheral: |*C*_*i*_| > 800, foveal: |*C*_*i*_| > 800, and organoid: |*C*_*i*_| > 400. The resulting percentage of cells in the low-expression category were – peripheral: 49.4%, foveal: 3.2%, and organoid: 0.5%. The majority of cells removed in the peripheral retinal samples were rods (estimated 77.7% of the 49.4%), which are over-represented in the peripheral data set and can dominate the analysis if not depleted (Macosko et al., 2015). The high-expression pool of each sample was embedded into a 2-dimensional matrix using the scVis method (Ding et al., 2018). Parameters supplied to scVis were – Adam optimizer, learning rate of 0.01, batch size of 512, *L*_2_ regularization strength of 0.001, perplexity of 10, and an ELU activation layer. Library normalization and transcript embedding were performed in Python 2.7 (NumPy v1.14.3, SciPy v0.18.1, scikit-learn v0.19.1, TensorFlow v1.9.0).

##### Infomap clustering and cluster merging

From the scores of cells for the first 100 principal components a graph was constructed using *K* nearest-neighbor search with *K* = 50 (LargeVis v0.2.1) (Tang et al., 2016). Clusters of cells with similar transcriptomes were identified by applying Infomap clustering (Rosvall and Bergstrom, 2008) to the graph (igraph v1.1.2). For each transcript, the random variable ‘cluster identity’ *T* was generated by mapping the cell identity *C* to its corresponding Infomap cluster *T*_*k*_. The values of T are the mutually disjoint subsets *T*_1_, *T*_2_, …, *T*_*n*_clust__ where *n*_clust_ is the total number of clusters. *T*_*k*_ represents a particular cluster of cells and contains the transcripts of each cell in that cluster. If |*T*_*k*_| is the number of transcripts in cluster *T_k_* then 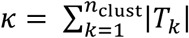. Similarly, if *m_T_k__* represents the number of cells in cluster *T*_*k*_ then 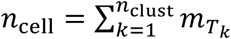. The contingency table between *T* and *G* is the count matrix **N** whose elements *N_kj_* are the number of transcripts observed in cluster *T*_*k*_ and gene *G*_*j*_. The normalized expression matrix 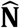 whose elements 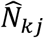 are the summed normalized expression values in cluster *T*_*k*_ and gene *G*_*j*_ generated by summing the normalized expression vectors 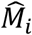 for each cell *C*_*i*_ in cluster *T*_*k*_. These clusters contain groups of cells with different transcriptomic “cell types”.

To assess cluster quality of each data set, we iteratively repeated the clustering 50 times. In each iteration, 85% of the high-expression cells were sub-sampled and the graph construction and Infomap clustering were repeated. Measures of cluster purity and stability during resampling were the same as those defined in similar studies to improve comparability (Shekhar et al., 2016).

Since the presence of batch effects in the scVis map (Supplemental Figure 5) indicated the potential for over-clustering, highly similar clusters were merged iteratively: pairs of clusters with a Pearson correlation coefficient of cluster-averaged gene expression *R*_P_ were merged if *R*_P_ > *θ*_merge_; *θ*_merge_ was initialized to 0.99 and clusters were iteratively combined for *R*_P_ > 0.90. High-expression cells were selected from the count matrix, and genes were selected if their mean-normalized log-range, mnr_*G_j_*_, was greater than the empirically estimated threshold 4.0.

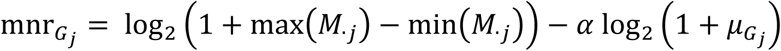

where max(*M*_·*j*_) and min(*M*_·*j*_) were the highest and lowest expression counts for gene *G*_*j*_ observed in the cell sample and *α* = 0.8 was a trend adjustment empirically estimated on held-out data. The number of genes thus selected were – peripheral: 632, foveal: 1273, and organoid: 705. Mergers were resolved in an inclusive fashion (e.g., if cluster 2 met the merger criterion with clusters 1 and 3, all three clusters would be merged together even if clusters 1 and 3 did not meet the merger criterion with each other).

##### Heatmaps of organoid single cell data

Let **S** represent the *n*_cell_ × 2 dimensional scVis map of organoid cell transcriptomes. The estimator 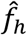 is a 2D Gaussian kernel density estimator used to calculate **Ŝ**, the estimated density of **S**. The kernel bandwidth *h* was estimated by Scott’s rule (Scott, 2015).

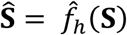

The elements of **Ŝ** are *Ŝ*_*xy*_ and its dimensionality was 341 × 341, evaluated between −17.0 and +17.0 with a step size of 0.1. For visualization purposes, an isodensity line was drawn at **Ŝ** = 2 × 10^−4^. A subset of organoid cells *C*^′^ ⊆ *C* was selected to analyze organoids from different ages, batches, etc. Letting 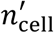 represent the number of cells in *C*’, the scVis map for these points is the matrix **S**^′^ with dimensions 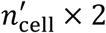. Holding *h* constant, the density of the subset was estimated as 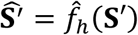 whose elements 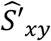 were evaluated at the same 341 × 341 locations.

The relative density of the subset of cells across the map is

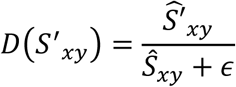

where the constant *ε* = 0.01 suppresses noise in regions with low density.

We used the weighted kernel density estimator *ĝ*_*h*_(**S**, *w*) to find **ψ**(*G*_*j*_), the estimated location for transcripts of gene *G*_*j*_ within the scVis map **S** (Supplemental Figure 3). In addition to the arguments accepted by 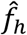, the estimator *ĝ*_*h*_ accepts the weight vector *w* which was set to the transcript count for gene *G*_*j*_ across all cells, *w* = *M*_·*j*_. The bandwidth *h* was held constant.

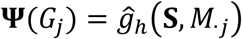

If the elements of **ψ**(*G*_*j*_) are *ψ*_*xy*_(*G*_*j*_), the relative density of transcripts across the map is

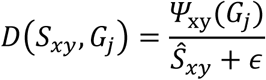

##### Stabilization of developing transcriptomes

Organoid transcriptomes at different ages were compared in the context of their scVis map kernel-density estimates. The 2-dimensional density estimates for a subset of cells 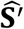 were *L*_1_ normalized by the total estimated kernel density 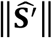 to create the probability mass function 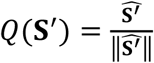. Two cell subsets, 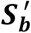 and 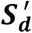, were compared by calculating their respective probability mass functions, 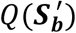 and 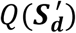, and calculating the Jensen-Shannon divergence between the two distributions. Briefly, if the average of their two probability mass functions is 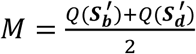 their Jensen-Shannon divergence is 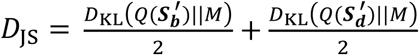 where *D*_KL_ is the Kullback-Leibler divergence.

##### Comparing the rate of organoid and human development

We used an available developing retina transcriptome from bulk (i.e. not single-cell) samples of human retina (Hoshino et al., 2017). Replicates were averaged and, to keep the approach region-agnostic, transcriptomes from specific retinal regions (e.g. peripheral retina) were excluded. Pseudo-bulk gene expression vectors were generated from our single cell data by using 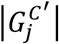, the number of unique transcripts observed for gene *G*_*j*_ within a subset of cells *C*’ (here, the subset of cells from an individual organoid). To focus subsequent analysis steps on genes likely to be modulated during development, we selected a gene *G*_*j*_ for analysis if the median-normalized range, dnr_*G_j_*_, of its vector of bulk/pseudo-bulk expression across tissues from different ages *B*_*G_j_*_ was greater than 0.5

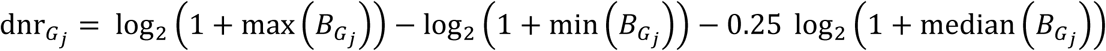

The number of genes thus selected was – organoids: 613; developing retina: 212. Because not all cell classes/types were present in each tissue (e.g., organoids lacked vasculature and immune cells, and the retinas lacked pigment epithelium) the intersection of the two gene sets, containing 93 genes, was used for both data sets.

A correlation matrix was constructed by determining the Spearman correlation coefficients, *R*_S_, of each individual retina/organoid to the others. To predict retina-equivalent ages of organoid transcriptomes we used ordinary least squares linear regression. Since genes outnumber the available retinas, we combatted overfitting by regressing against a transcriptome’s correlations to the retina transcriptomes. First, the model was trained to predict retinal ages from retina-retina correlations. The training data was the set of vectors of *R*_S_ between each retinal transcriptome and all retinal transcriptomes, and the target values were the retinal ages. To evaluate the model, leave-one-out cross validation was applied; the age of each retinal transcriptome was predicted by a model trained on the other retinal transcriptomes. Second, the trained model was applied to the retina-organoid data; the input for an organoid was the vector of *R*_S_ between its transcriptome and each retinal transcriptome. Because the pattern of positive correlations extended for weeks in either direction, the model predicted a retinal-equivalent age outside of the range of retina ages for some organoid transcriptomes.

##### Multiplet removal

Multiplets occur when one cell barcode shares transcripts from multiple single cells due to incomplete dissociation or well-dissociated single cells being placed in the same droplet. Transcriptomes likely to be multiplets were identified by the presence of genetic markers for two or more cell classes or sets of classes. The sets of markers were – rods: *GNGT1*, *NRL*, *PDE6G*, *RHO*; cones: *ARR3*, *CNGA3*, *OPN1LW*, *OPN1MW*, *OPN1SW*, *PDE6H*; horizontal cells: *LHX1*, *LNP1*, *ONECUT1*, *VAT1L*; bipolar cells: *GRIK1*, *IRX6*, *LRTM1*, *PCP2*, *PRKCA*, *TRPM1*, *VSX1*, *VSX2*; amacrine cells: *GAD1*, *SLC6A9*, *TFAP2A*, *TFAP2B*; ganglion cells: *POU4F2*, *NEFL*, *NEFM*, *RBPMS*, *SLC17A6*, *SNCG*, *THY1*; pigmented cells: *BEST1*, *MITF*, *MLANA*, *TJP1*, *RPE65*; glial cells: *CRABP1*, *GFAP*, *GLUL*; endothelial cells, mural cells and fibroblasts: *ACTA2*, *COL1A2*, *EGFL7*, *PDGFRB*, *PROCR*, *VWF*; immune cells: *AIF1*, *CD2*, *CD48*, *CX3CR1*, *HBB*, *IL32*, *JCHAIN*, *LST1*.

For a cell *C*_*i*_ and a set of marker genes *G*′ containing gene indices J, a score was created by summing transcript counts, score 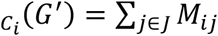. If marker specificity for the target cell class was perfect and background RNA was absent, the score distribution would contain a mode with positive expression (cells of the class) and a non-overlapping mode at zero (cells not of the class). In practice, the specificity is imperfect and the distribution more complicated. The threshold score for separating cells of the class from cells not of the class were empirically estimated for each set of marker genes independently. Cells with a score_*Ci*_ above threshold for two or more marker sets were flagged as multiplets.

Because this process followed clustering and embedding, it was possible to validate thresholds by visualizing multiplets on the scVis map. The transcriptomes of cells removed as multiplets were commonly present in low density regions of scVis maps, especially in the space between pairs of populous cell types (e.g., between rods and rod bipolar cells, rods and Müller cells) which is a rational placement for doublets (Supplemental Figure 5). Because score_*Ci*_ relies on a small number of genes and transcripts are prone to dropout it is susceptible to false negatives. Therefore, cells were also labelled multiplets if they shared a cluster with a high percentage of multiplets: > 30% for adult retina; > 15% for organoid. The total percentages of cells removed as multiplets were: peripheral: 5.7%; foveal: 6.1%; developing organoids: 1.0%; developed organoid: 5.0%.

##### Cell type composition of single-cell sequencing data

Since the cell-class composition in the Infomap clusters was biased by the removal of low-expressing cells, the marker gene sets were used to estimate the cell-class composition of single-cell transcriptomes. Using the same threshold scores set when finding multiplets, each cell was assigned a cell class if it was above the threshold score for that class. No cell was above threshold for more than one class, as these were removed as multiplets. The number of cells not above threshold for any class was 4.6% in the fovea retina, 10.4% in the peripheral retina, and 30.0% in the developed organoid. For each cell class, the compositions in two data sets were compared by forming a contingency table for cells inside/outside of the class in the foveal-retina/peripheral-retina/developed-organoid and performing a χ^2^ test.

##### Regional transcriptomic character of cell classes

The following process was repeated for different cell classes or types. If multiple clusters belonged to a single cell class, those clusters were combined and analyzed as a single cluster. Recall from above that *T* is the cluster identity of transcripts, and the subset of transcripts belonging to a cluster is *T*_*k*_. The number of cells in cluster *T*_*k*_ is *m*_*T_k_*_ and 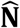 is the expression matrix of size *n*_clust_ × *n*_gene_ whose elements 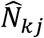 are the sum of the normalized expression values in cells of cluster *T*_*k*_ for gene *G*_*j*_. Since peripheral and foveal data sets were processed separately, superscripts fov and per are employed to distinguish between variables referring to the two data sets. Cells from foveal retina and developed organoid were scaled to have the same library size as the peripheral retina. To simplify the notation, the same index *k* is used to refer to the cluster in both data sets, as if the cluster indices were ordered by cell class. For example, the notation for cluster *T_k_* in the peripheral and foveal retina are 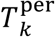 and 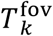 and for normalized expression matrix 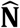 they are 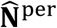 and 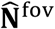. The cell-averaged expression of gene *G*_*j*_ within cluster *T*_*k*_ in the foveal retina was

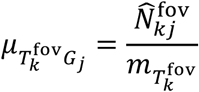

And the cell-averaged mean expression for the two regions is

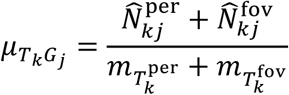

In addition to considering *μ*_*T_k_G_j_*_, the average transcript counts within cluster *T*_*k*_, we also considered *μ*_*T_o_G_j_*_, the average transcript counts in all clusters other than cluster *T*_*k*_

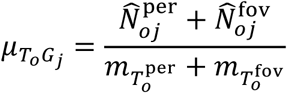

50% of the single cells were selected at random and held-out for later cross-validation. On the remaining transcriptomes, genes were classified as overexpressed if: 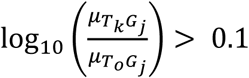, *μ*_*T_k_G_j_*_ > 1 and *μ*_*T_o_G_j_*_ < 10. To limit the potential influence of low levels of background transcripts from other clusters, further analysis of the cluster’s transcriptome was restricted to these overexpressed genes.

For gene *G*_*j*_ in the context of cluster *T*_*k*_, the index of regional specificity was

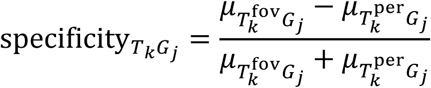

The significance of regional expression differences for gene *G*_*j*_ was evaluated by the Mann-Whitney U test with continuity correction on the normalized expression vectors 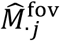 and 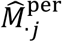. Since *P*-values falling below 1.8×10^-307^ underflowed to zero, they were visualized by assigning a random value between 10^-350^ and 10^-400^.

A classifier was trained by least squares regression to predict the region-of-origin for single cells of the cluster based on their transcriptomes. The combined foveal and peripheral data sets were standardized and each set was shuffled and trimmed so that the number of cells per region, *m*, was equal and at most 1000. The peripheral and foveal transcriptomes were combined to create a 2*m* × *d* set of observations to train the model, where *d* is the number of overexpressed genes for that cluster. The model output to be predicted was the length 2*m* vector containing the target regional labels from paired observations; the label for peripheral retina was mapped to −1, foveal retina to +1. For cell *C*_*i*_, the model for the regional classifier was

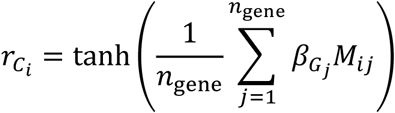

The model coefficients *β* are an *n*_gene_ length vector whose element *β*_*G_j_*_ is initialized to 10% of the regional specificity index for gene *G*_*j*_. The optimization cost function was least squares with an *L*_2_ regularization weight of 1×10^-3^. After fitting the model, held-out data was used for cross-validation. Let 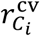 be the regional classifier score for cell *C*_*i*_ from the cross validation with held-out data. Let *y*_*Ci*_ be the target regional labels of the cross-validation and *n*_cv_ be the number of cells in the cross-validation. The classifier’s coefficient of determination, *R*^2^, was then

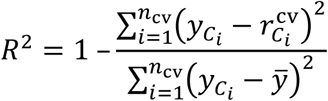

where

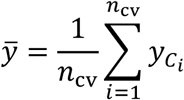

For each cell class that was present in organoids, cell transcriptomes underwent the same pre-processing and their *r*_*Ci*_ was calculated to predict their regional character. Organoids cells were defined as “overlapping” the adult cell distribution if they were below the 95^th^ percentile of peripheral cells or above the 5^th^ percentile of foveal cells. If a mutually overlapping interval was present, cells in its range were excluded.

##### Closeness of organoids to adult cell classes

The procedure below was applied to a cell class. Genes differentially expressed for a cell class were defined by the metric

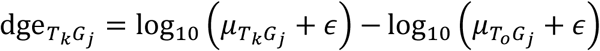

where *ε* = 0.5 was a constant. Principal component analysis was performed using the 20 genes with the highest dge_*T_k_G_j_*_ as the dimensions, and a combination of (i) all peripheral retinal cells from the cell class and (ii) a number of organoid cells, equal to the number of peripheral retinal cells, subsampled without respect to organoid age or cell class. The resulting principal components were used to project the remaining organoid cells.

The distribution of the first two principal components (PC1, PC2) of adult cells was fit using the graphical lasso algorithm. The Mahalanobis distance of each organoid cell to this distribution was then calculated to assess how “close” organoid cells were to the adult cell distribution within the space of the top cell class-specifying genes.

##### Mapping genes to cell classes

The distribution of genes across clusters was determined by taking µ_*T_k_G_j_*_, the average library-normalized expression of gene *G*_*j*_ within each cluster *T*_*k*_, and dividing by ‖µ_·*G_j_*_‖ the sum of the average expression vector across all clusters (as above, we are using the *L*_1_ norm)

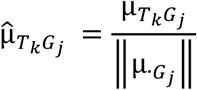

##### Cell type specificity of genes

Relationships between gene and cluster identity were assessed probabilistically. Recall from above that **N** is the count matrix whose elements *N*_*kj*_ are the number of transcripts observed in cluster *T*_*k*_ and gene *G*_*j*_ and that *κ* is the total number of transcripts. The probability of observing a specific combination of cluster *T*_*k*_ and gene *G*_*j*_ is the joint categorical distribution

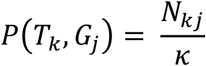

such that

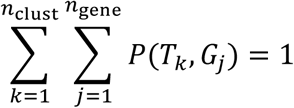

The two marginal distributions are 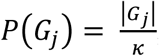, representing the probability of observing gene *G_j_*, and 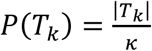, the probability of observing cluster *T_k_*. The entropy for a given clustering *T* is

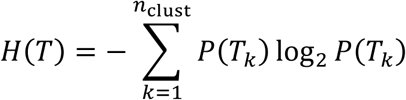

The gene entropy is given by

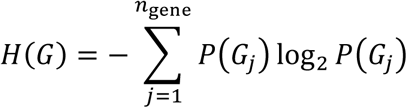

The entropy for the joint distribution is

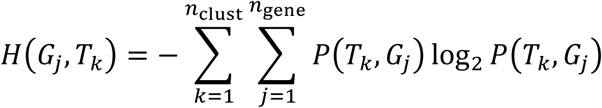

We utilize these entropies to assess the cluster specificity of a gene – whether transcripts of the gene are concentrated within some clusters and not others. An entropy of zero means there is no uncertainty (i.e., 100% probability of the gene and cluster identities coinciding) and entropy is at its maximum for perfectly random probability distributions (i.e., equal probability of all gene and cluster combinations). For a transcript from a given gene *G*_*j*_, the uncertainty about the cluster identity is the conditional entropy

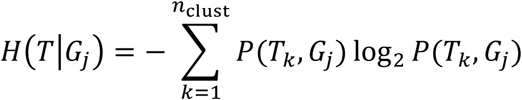

The specificity of individual disease-associated genes spg_*G_j_*_ was defined relative to the normalized entropy

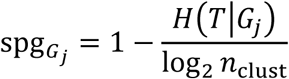

The significance of spg_G_j__ was assessed using a Monte Carlo permutation test. The following process was repeated a total of *n*_iter_ = 1,000 times: (i) *C*, the cell identity of transcripts, was randomly shuffled to create *C*’; meanwhile, its paired observation *G* was held constant; (ii) the existing Infomap clustering was applied to *C*’ to generate *T*’, the cluster identity of the cell-shuffled transcripts; thus, the cell type identities are randomized at the transcript level without changing *P*(*G*_*j*_) or *P*(*T*_*k*_); (iii) the count matrix with shuffled cell-type identities 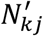 was generated from *T*’ and *G*; (iv) the conditional entropy of gene *G*_*j*_ was determined for the permuted clustering *H*(*T*′|*G*_*j*_) and the specificity 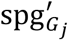 calculated; (v) the value of 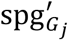 was stored and if it was greater than the specificity in the non-permuted clustering spg_*G_j_*_ a counter *b* was incremented.

Once concluded, if *b* > 10 then 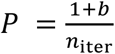, otherwise the parameters *ϕ* of a generalized Pareto distribution *F*_*ϕ*_(*z*) were fit to the exceedances, *z*, which are the permutation test occurrences of 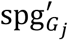 that exceeded the threshold *φ* (Knijnenburg et al., 2009). The initial value of *φ* was set such that there were *n*_exceed_ = 250 exceedances, and the goodness-of-fit for the generalized Pareto distribution was evaluated by the Kolmogorov-Smirnov test. If the fit was poor (null hypothesis that distributions are identical rejected, P < 0.05), *φ* was adjusted upward such that *n*_exceed_ was lowered in steps of 10 until the fit was good. The test statistic spg_*G_j_*_ was then evaluated on the fit distribution to generate the P-value

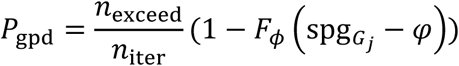

It was also useful to assess the comodulation of a subset of genes, *G*^′^ across the set of clusters. This method was used to quantify the relationship between gene identity and cluster identity for (i) a set of genes marking different cell types (marker-defined cell types) and (ii) the subset of genes associated with a disease. To do so, we calculated the mutual information between the cluster identities, *T*, and a subset of gene identities, *G*′

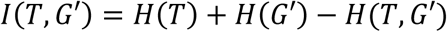

If the mutual information is not zero, then knowledge of a transcript’s genetic identity reduces the uncertainty about its cluster identity. The reverse is also true, but we focus on the case where genes are given. The cluster identity information gained by knowing the gene identity expressed relative to the uncertainty about cluster identity is the uncertainty coefficient

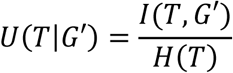

the *P*-value of which was estimated by the Monte Carlo permutation test described above with one modification: in steps (iv) and (v) the statistic being tested was the uncertainty coefficient rather than the specificity.

##### Disease genes

The list of disease genes was drawn from the NIH Genetics Home Reference (https://ghr.nlm.nih.gov). We included all genes they listed as being associated with a disease as of August 13^th^ 2018. To ensure an unbiased representation, the list was not revised based on our own observations. In the disease maps, four disease genes were not visualized because they were not expressed in at least one data set (age-related macular degeneration-associated genes: *F13B*, *CFHR2*, *CFHR3*, and *CFHR5*).

##### Development of workflow for transcriptome analysis

The transcriptome analysis presented here was developed in three stages. First, the workflow was designed and algorithms selected using transcriptomes from non-human primate retinas. Second, values presented as “empirically estimated” were manually set by an experimenter using data from the first human retinal donor and organoid data up to 24 months of age. Third, the full data set was analyzed. Few adjustments were made between the second and third steps; most notably the input space for Infomap clustering was changed from 1000 over-dispersed genes to 100 principal components.

## Supplemental Figure Legends

**Supplemental Figure 1.**
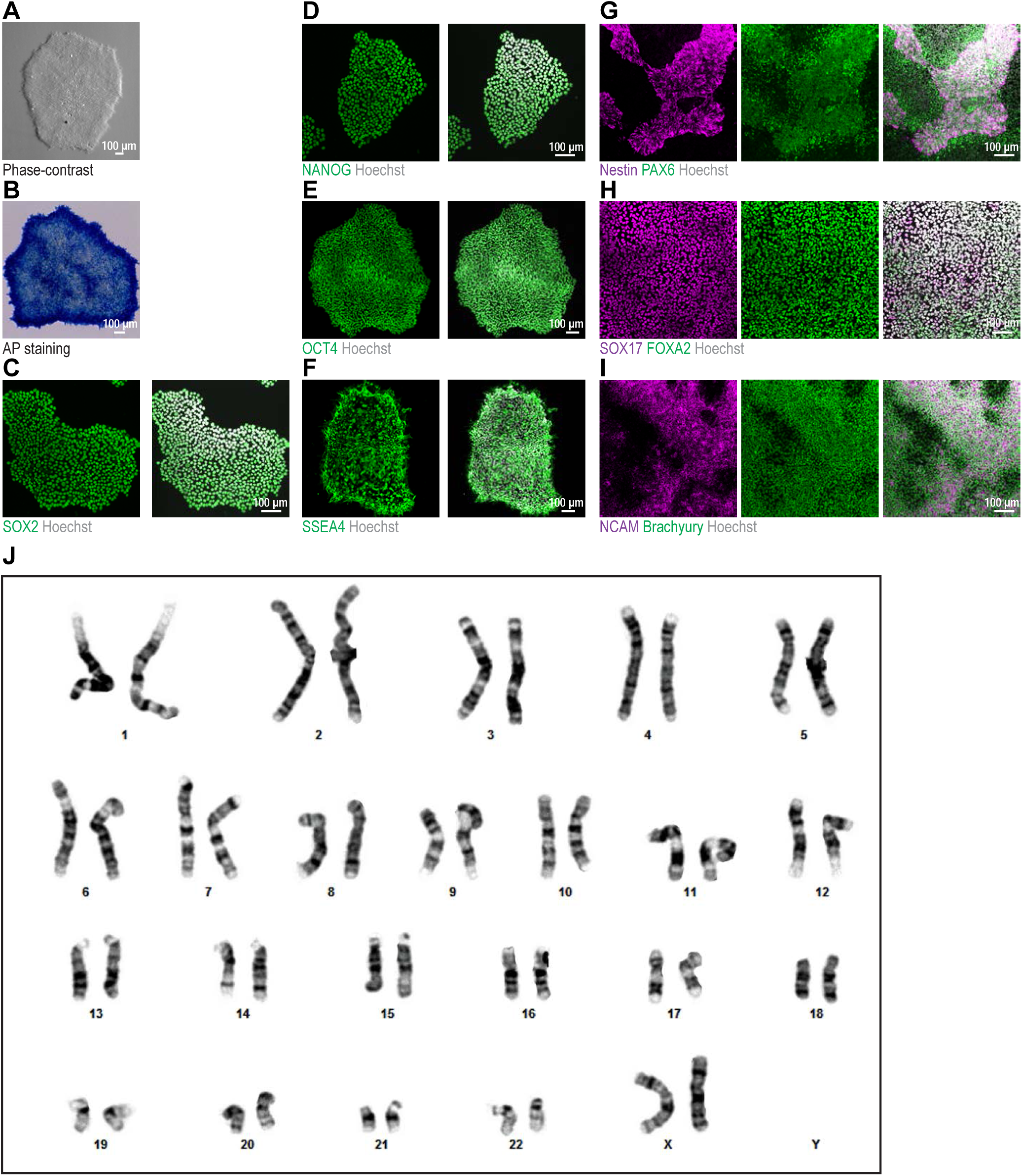
Characterization of F49B7 iPSCs. (A) Brightfield image of F49B7 iPSC colony. (B) Brightfield image of iPSC colony. Alkaline phosphatase stain (AP; pluripotency marker). (C – F) Confocal images of iPSCs. Green, antibody for pluripotency markers; white, Hoechst (nucleus marker). (C) SOX2, (D) NANOG, (E) OCT4 and (F) SSEA4. (G – I) Confocal images of iPSCs directly differentiated into the three germ layers. White, Hoechst. (G) Ectoderm; magenta, Nestin; green, PAX6 (ectoderm markers). (H) Endoderm; magenta, SOX17; green, FOXA2 (endoderm markers). (I) Mesoderm; magenta, NCAM; green, Brachyury (mesoderm markers). (J) G-banded karyotyping of F49B7 iPSCs.

**Supplemental Figure 2.**
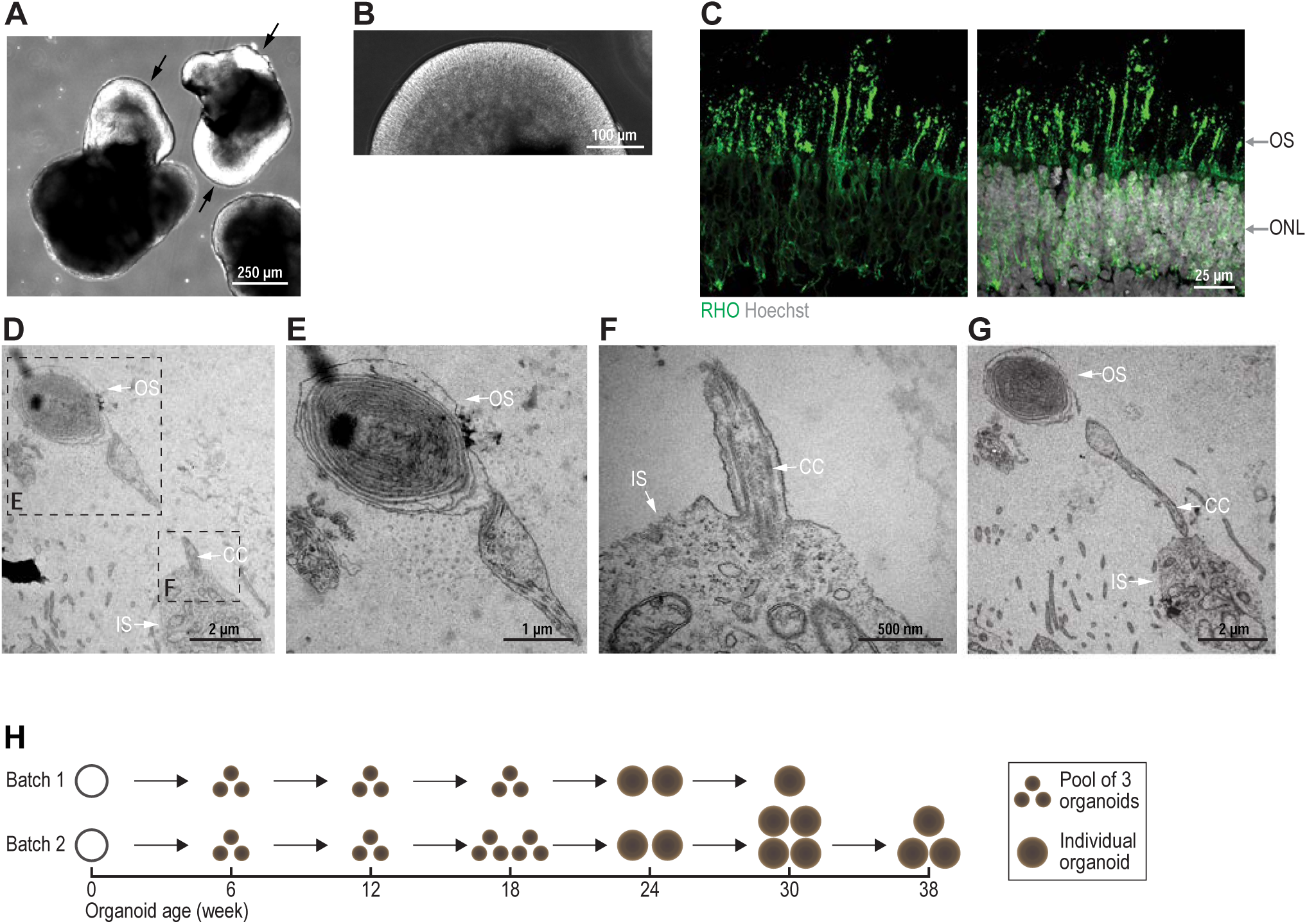
Organoid characterization. (A) Retinal neuroepithelium (arrows) on a five-week old organoid. (B) Retinal neuroepithelium appears phase-bright and columnar. (C) Confocal image. Green, RHO antibody (rod outer segment marker); white, Hoechst (nucleus marker). OS, outer segment; ONL, outer nuclear layer. (D – G) Electron microscope images of photoreceptor outer segment (OS; D, E, G), inner segment (IS; D, F, G) and connecting cilium (CC; D, F, G). D and G are projections from serial sections. (H) Scheme of the time course with which organoids were sampled for single-cell RNA sequencing.

**Supplemental Figure 3.**
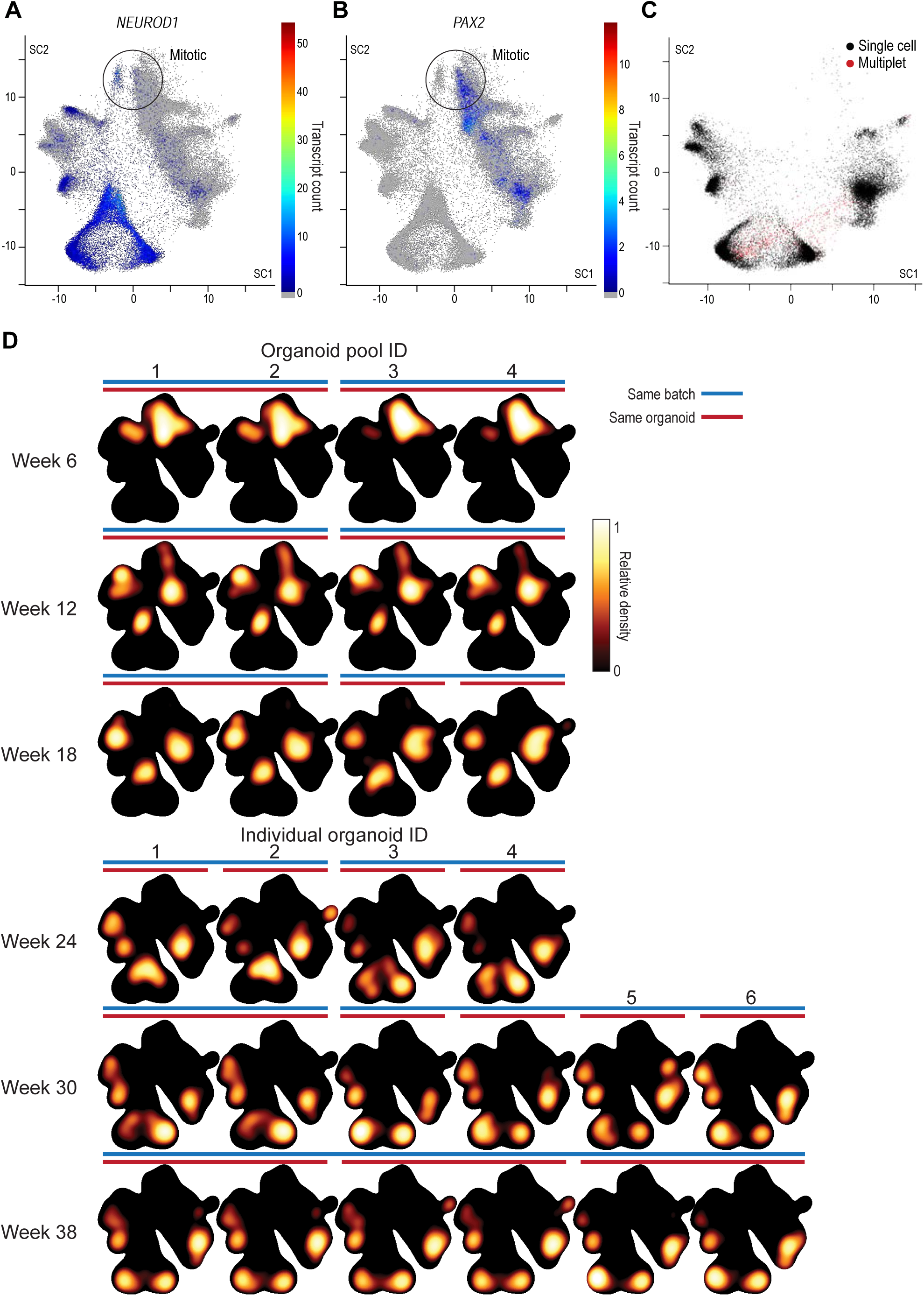
Reproducibility of organoids. Location of transcripts of (A) neurogenic and neuronal marker *NEUROD1* and (B) gliogenic marker *PAX2* in organoid scVis map. Color, transcript count; colormap at right. (C) Cells removed as multiplets (red) in developed organoids shown on the developed organoid scVis map. (D) Heatmap of relative cell density in 28 different samples with indicators for organoid age (row), pooling method (top, pool; bottom, individual), batch (blue bar), and replicates from the same organoid (red bar).

**Supplemental Figure 4.**
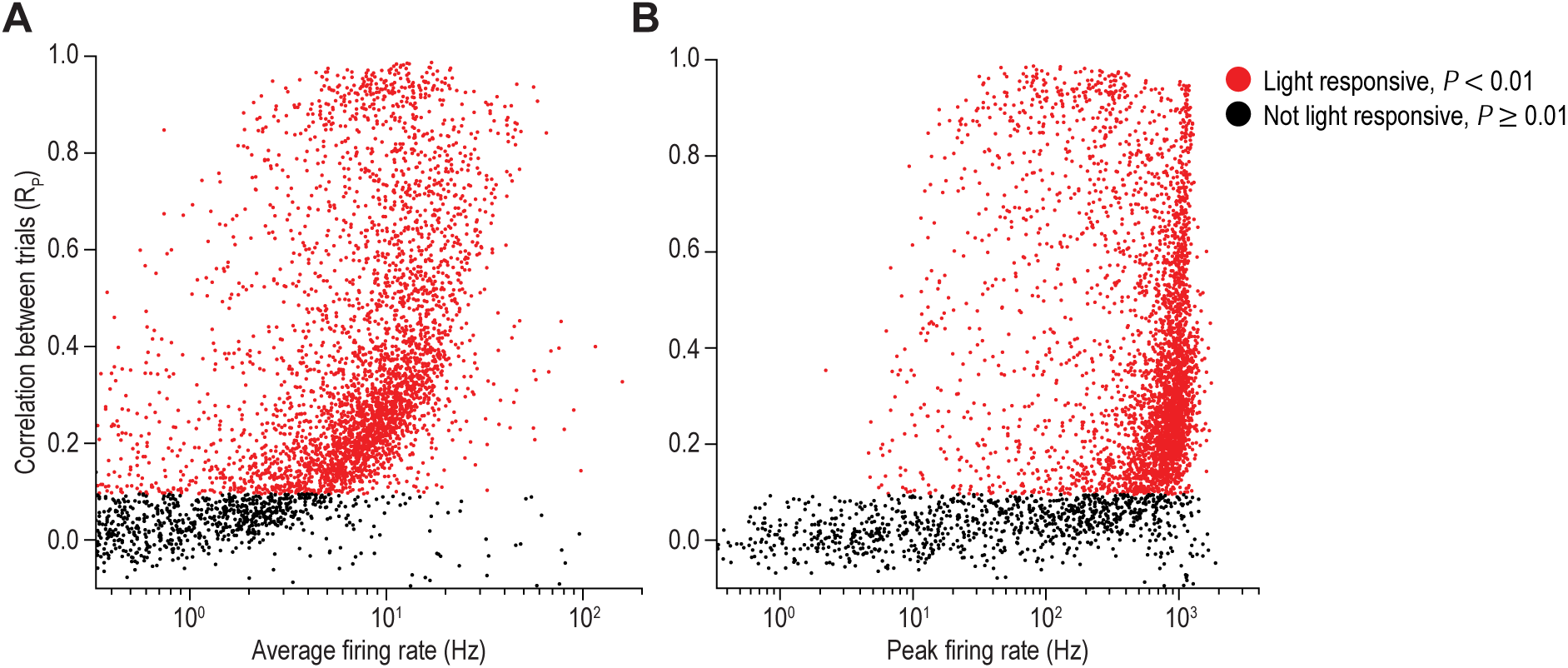
Firing rate statistics of adult retinal ganglion cells. Scatter plots comparing the correlation in ganglion cell firing rates across stimulus repetitions to their (A) average firing rate and (B) peak firing rate. Red, cells significantly light responsive (*P* < 0.01). Black, cells not light responsive (*P* ≥ 0.01)

**Supplemental Figure 5.**
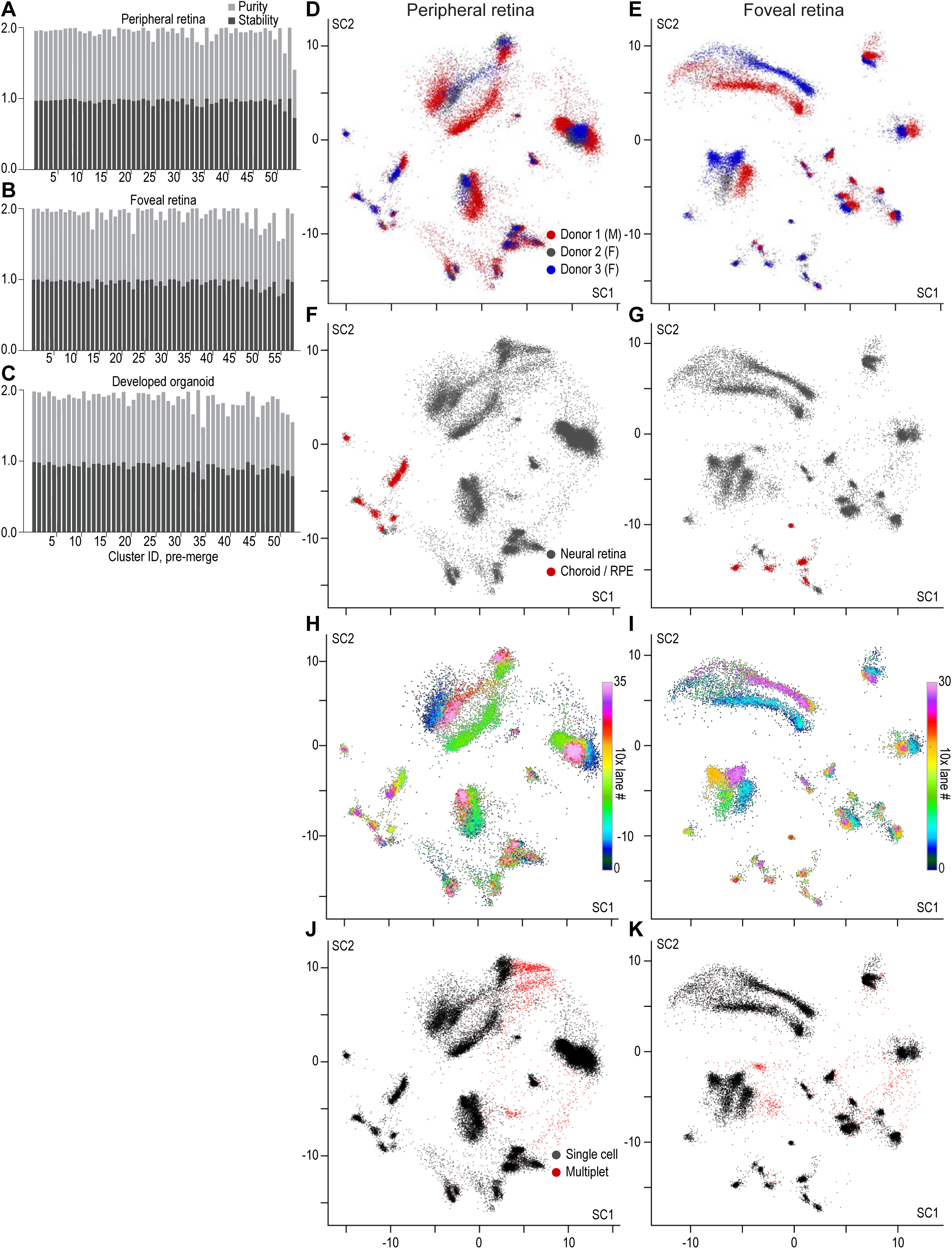
Infomap clustering quality and sources of variability in adult retinal scVis maps. Cluster quality was evaluated in a bootstrap process to evaluate the Infomap clusters of (A) peripheral retina, (B) foveal retina and (C) developed organoids. Light grey bars, cluster purity. Dark grey bars, cluster stability. (D – K) The peripheral (left) and foveal (right) scVis maps labelled according to (D – E) donor identity and sex; M, male; F, female, (F – G) tissue identity and (H – I) 10x Chromium lane. (J – K) Cells identified as likely multiplets and removed from the analysis.

**Supplemental Figure 6.**
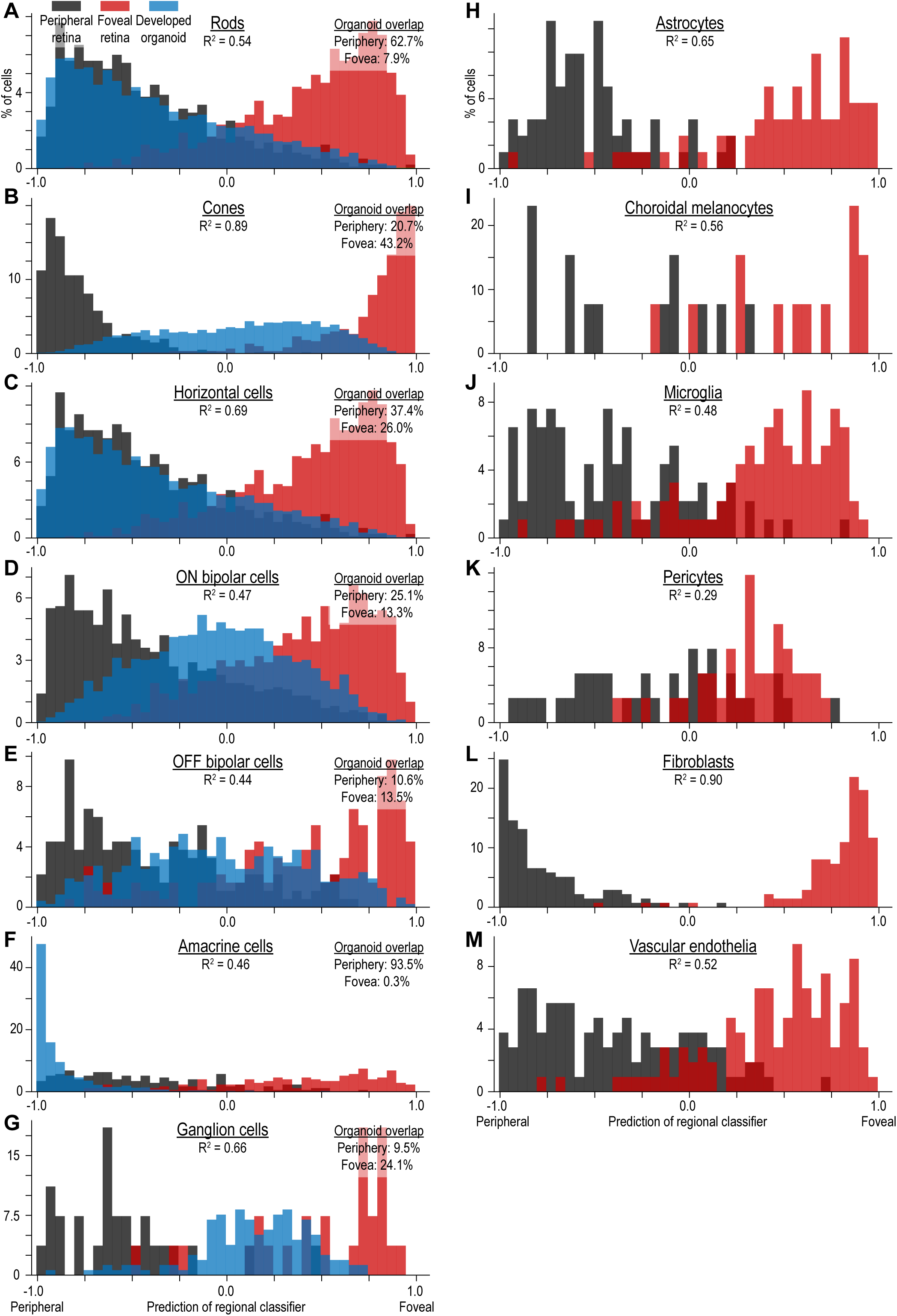
Regional character of cell types in adult retina and organoids. Predictions from a classifier trained to predict the region of origin (peripheral, −1.0; foveal, +1.0) of adult cells based on their transcriptome. Black bars, peripheral cells held out from training. Red bars, foveal cells held out from training. R^2^, coefficient of determination. (A – G) For cell classes present in organoids, the transcriptomes of organoid cells were regionally classified. Blue bars, organoid cells.

**Supplemental Figure 7.**
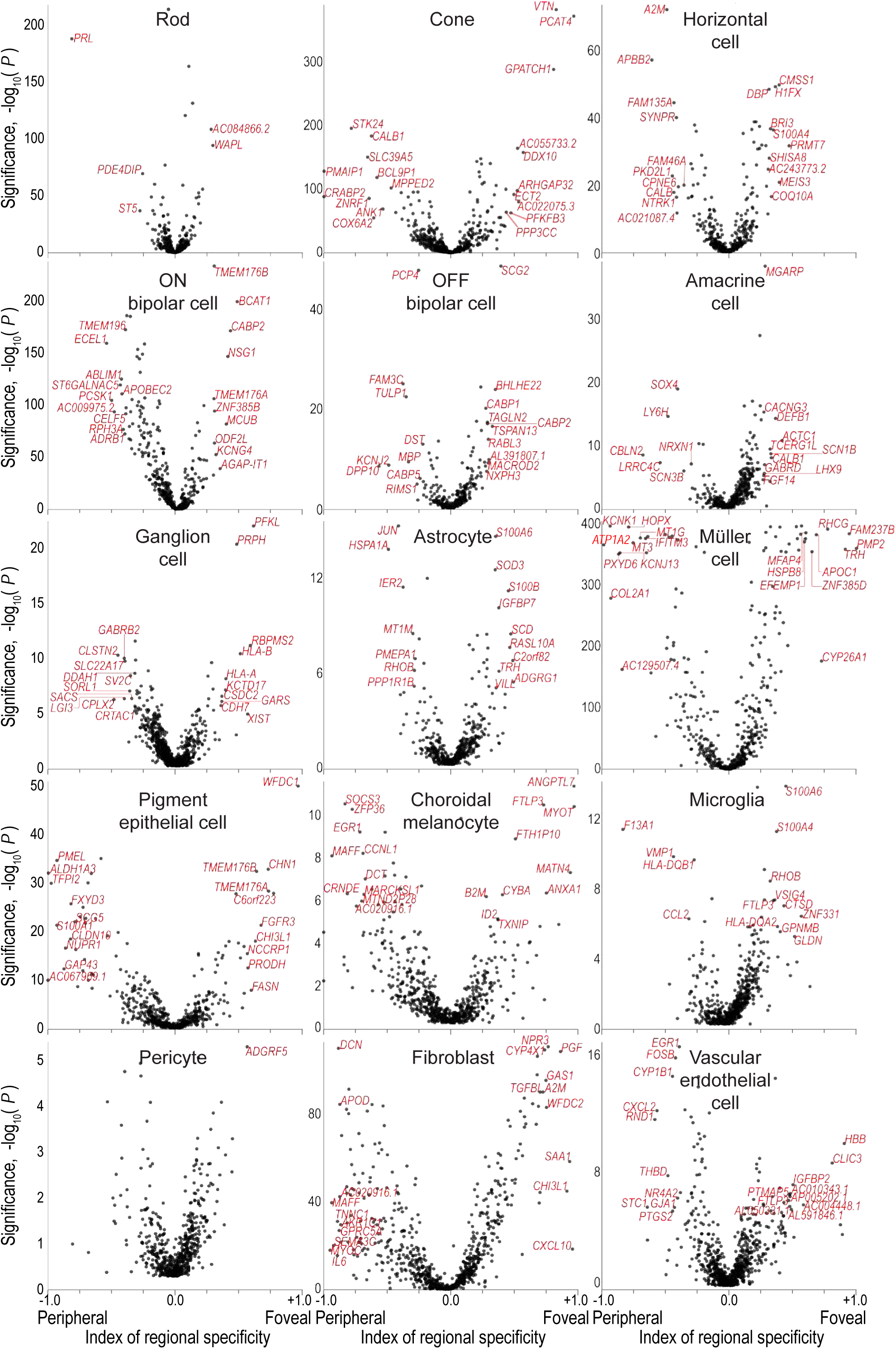
Regional differences in gene expression within cell types of the adult retina: fovea versus periphery. Regional specificity of genes for the peripheral (negative values) or the foveal retina (positive values) within different cell classes. Scatter plots, different cell classes. Black points, genes. Red labels, names of top 10 peripheral specific genes and top 10 foveal specific genes with a *P* < 10^-5^ and an absolute specificity > 0.25.

**Supplemental Figure 8.**
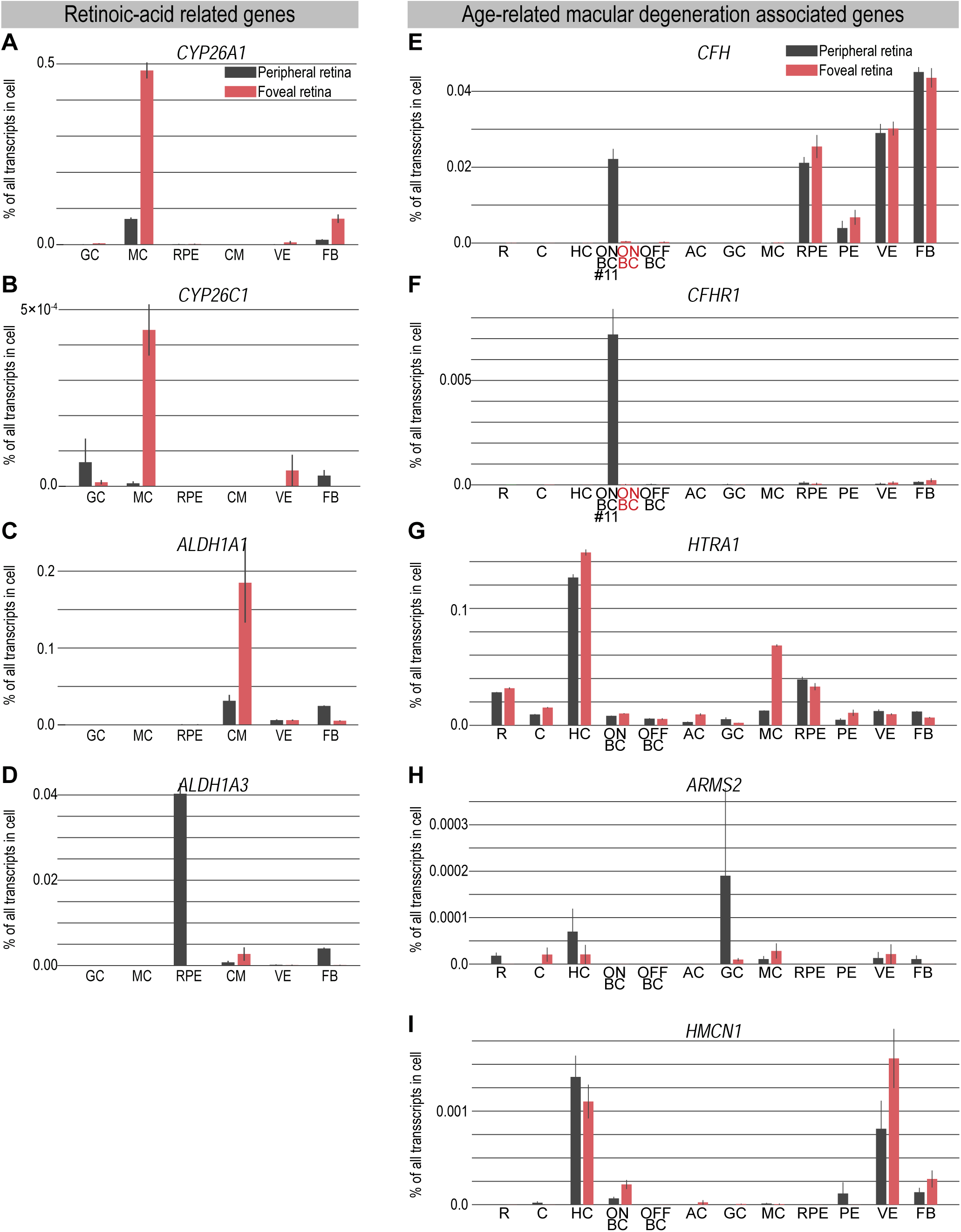
Cell type expression of retinoic-acid related and age-related macular degeneration associated genes. (A – I) Expression levels in selected cell types. Black bars, peripheral retina; red bars, foveal retina. Cell type acronyms are from Figure 5B. Error bars, ±1 SE. (A – D) Genes for retinoic acid synthesis (*ALDH1A1*, *ALDH1A3*) and catabolism (*CYP26A1*, *CYP26C1*). (E – I) Genes associated with age-related macular degeneration. (E-F) Peripheral Infomap cluster #11, a type of cone ON bipolar cell, is compared to all foveal ON bipolar cells.

**Supplemental Figure 9.**
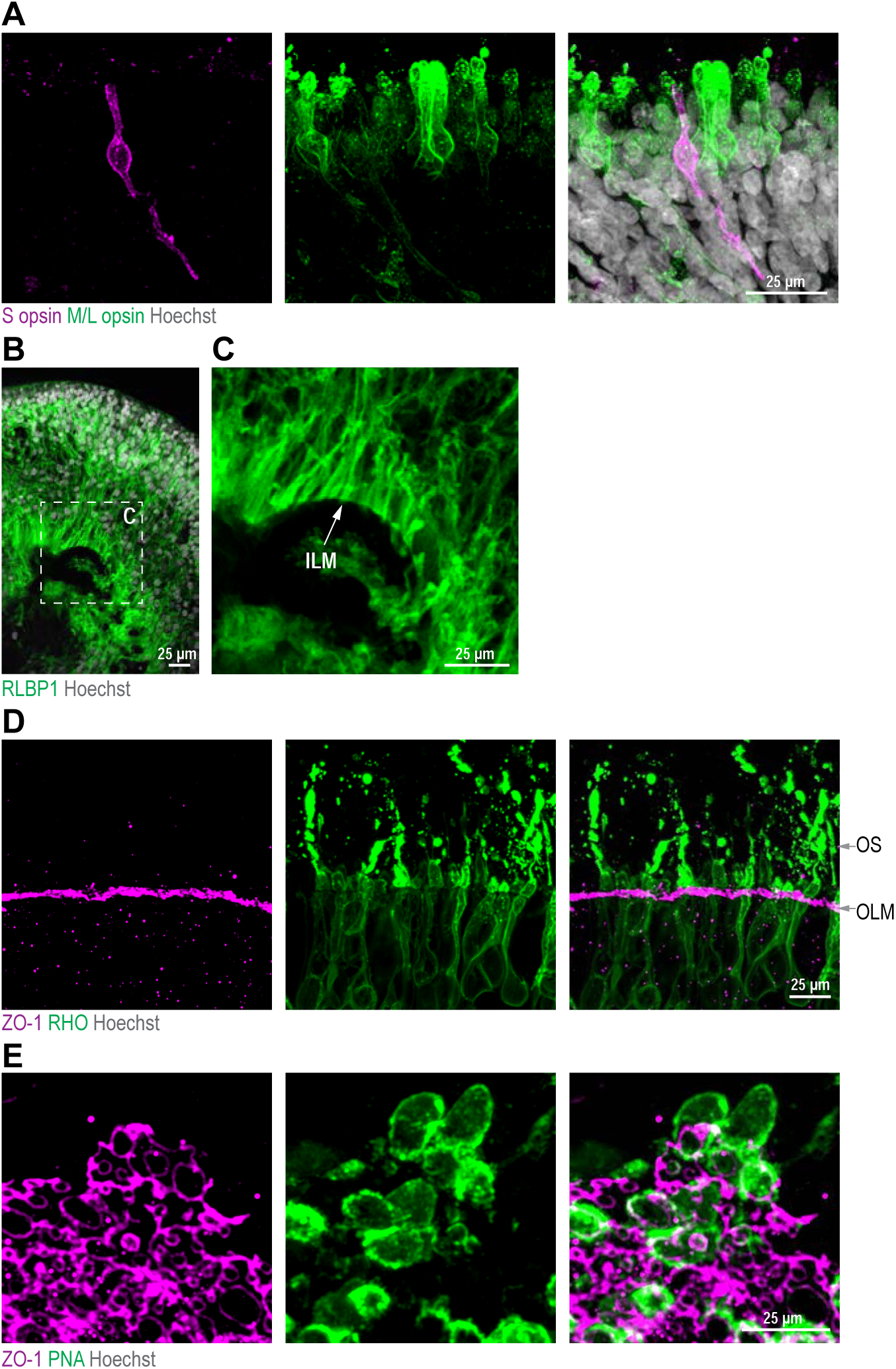
Cell types and structures in developed organoids. (A – E) Confocal images. (A) Magenta, S opsin antibody (blue cone marker); green, M/L opsin antibody (green/red cone marker); white, Hoechst (nucleus marker). (B) Green, RLBP1 antibody (Müller cell marker); white, Hoechst. (C) Boxed area from B. ILM, inner limiting membrane. (D) Magenta, ZO-1 antibody (outer limiting membrane marker); green, rhodopsin antibody (RHO, rod outer segment marker). OS, outer segment; OLM, outer limiting membrane. (E) Top view of ZO-1 rings, maximum intensity projection. Magenta, ZO-1 antibody; green, PNA (cone outer segment marker); white, Hoechst.

**Supplemental Figure 10.**
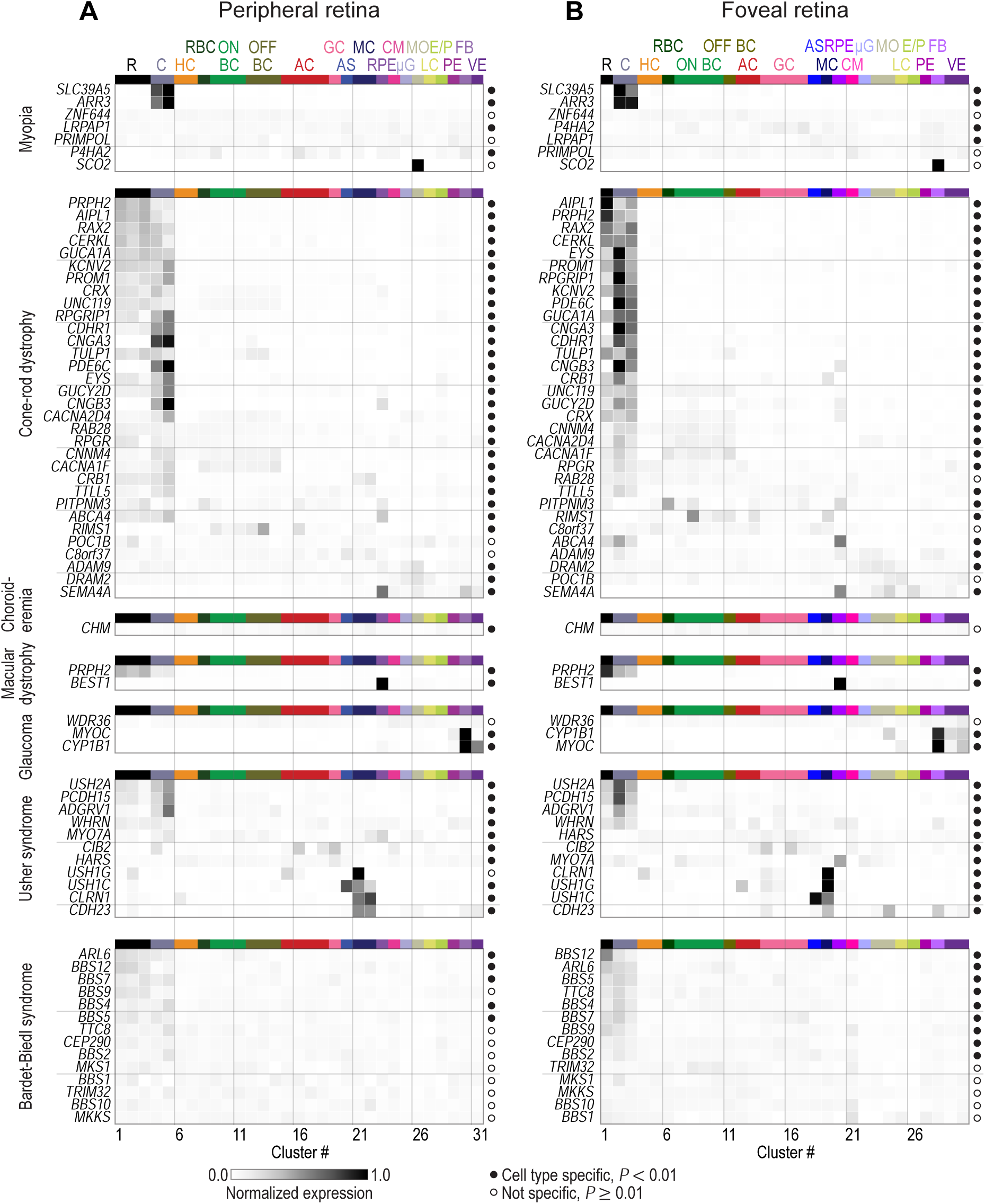
Supplementary disease map for the peripheral and foveal retina. Normalized expression of disease genes (rows) within cell types (columns) in (A) peripheral and (B) foveal retina. Left, names of diseases and associated genes. Colormap at bottom left of A, level of intra-gene normalized expression. Filled circle, gene significantly cell type specific (*P* < 0.01); empty circle, gene not cell type specific (*P* ≥ 0.01). Cell type colors and acronyms are according to Figure 5B.

**Supplemental Figure 11.**
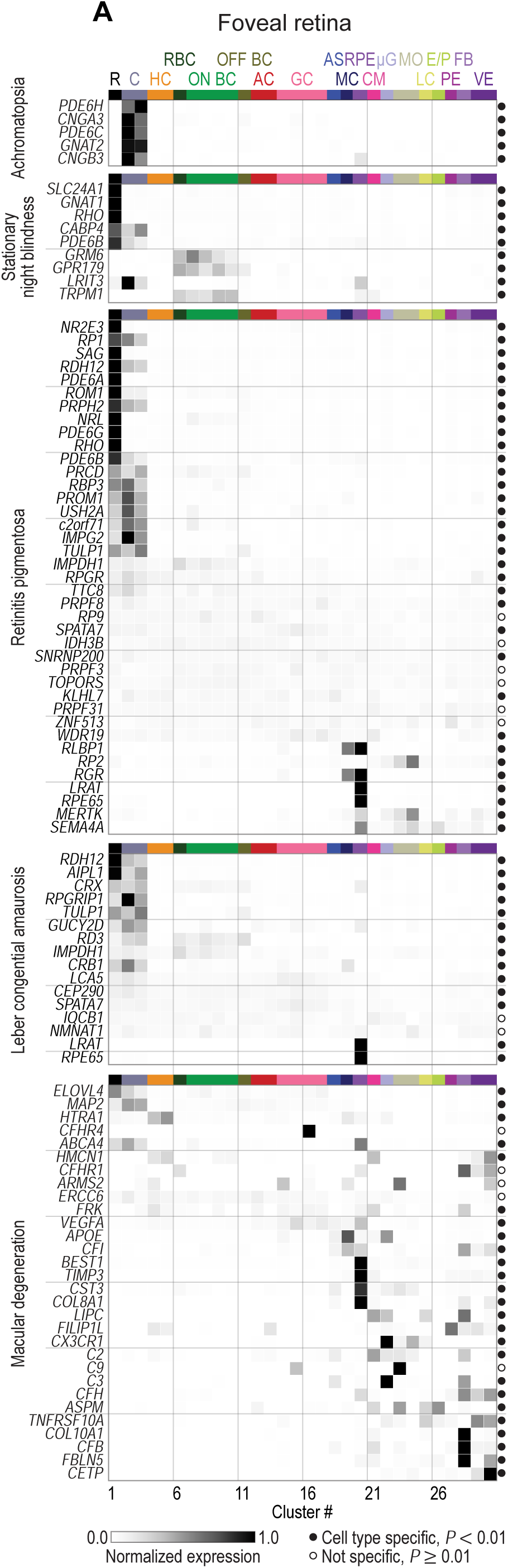
Disease map for foveal retina. (A) Normalized expression of disease genes (rows) within cell types (columns) in foveal retina. Left, names of diseases and associated genes. Colormap at bottom left, level of intra-gene normalized expression. Filled circle, gene significantly cell type specific (*P* < 0.01); empty circle, gene not cell type specific (*P* ≥ 0.01). Cell type colors and acronyms are according to Figure 5B.

**Supplemental Figure 12.**
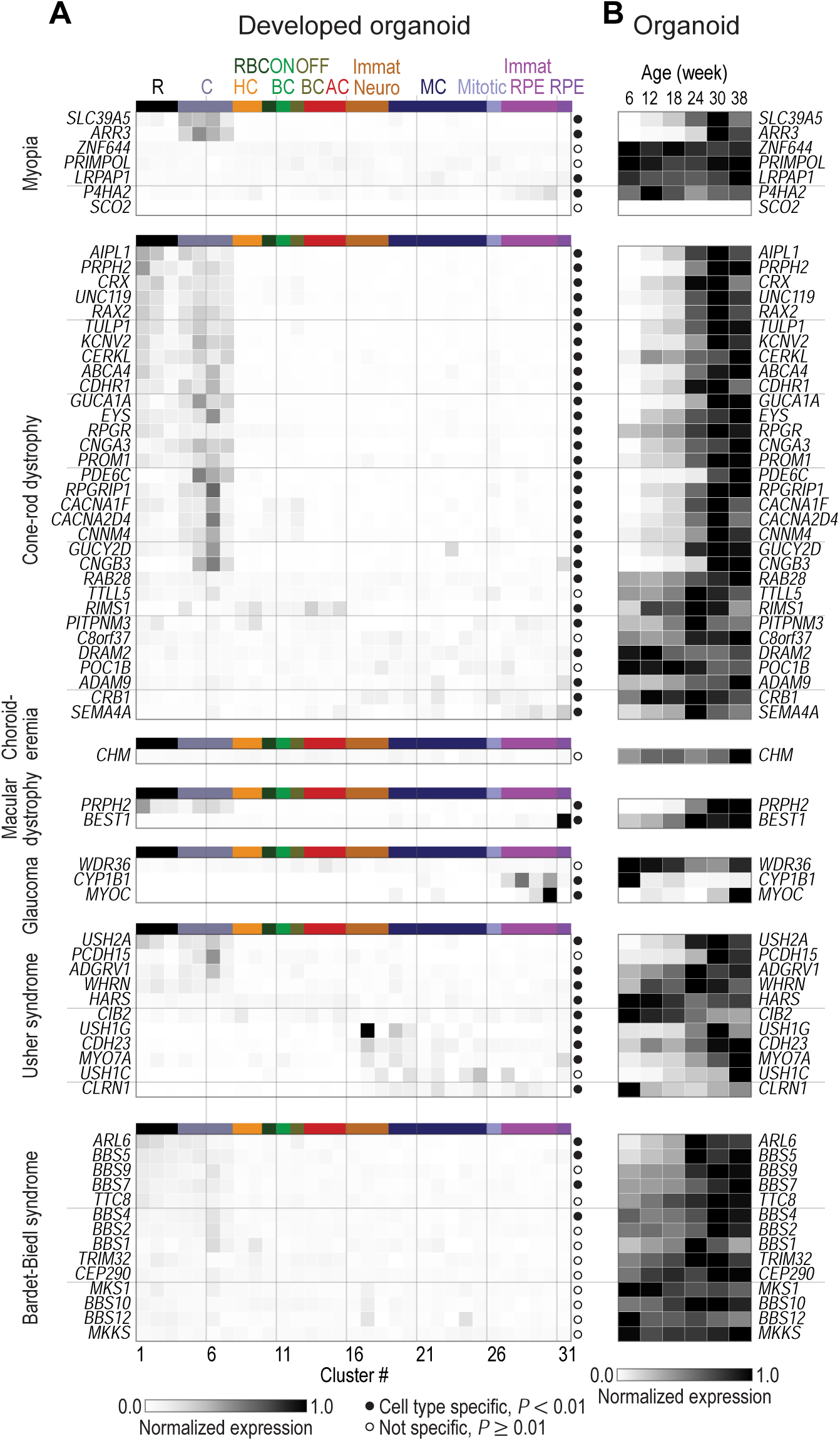
Supplementary disease map for developed organoids. Normalized expression of disease genes (rows) within cell types (columns) in developed organoid. Left, names of diseases and associated genes. Colormap at bottom left of A, level of intra-gene normalized expression. Filled circle, gene significantly cell type specific (*P* < 0.01); empty circle, gene not cell type specific (*P* ≥ 0.01). Cell type colors and acronyms are according to Figure 5B. (B) Age (columns) dependence of disease gene (rows) expression within organoids. Colormap at bottom of B, level of min-max normalized expression.

**Supplemental Figure 13.**
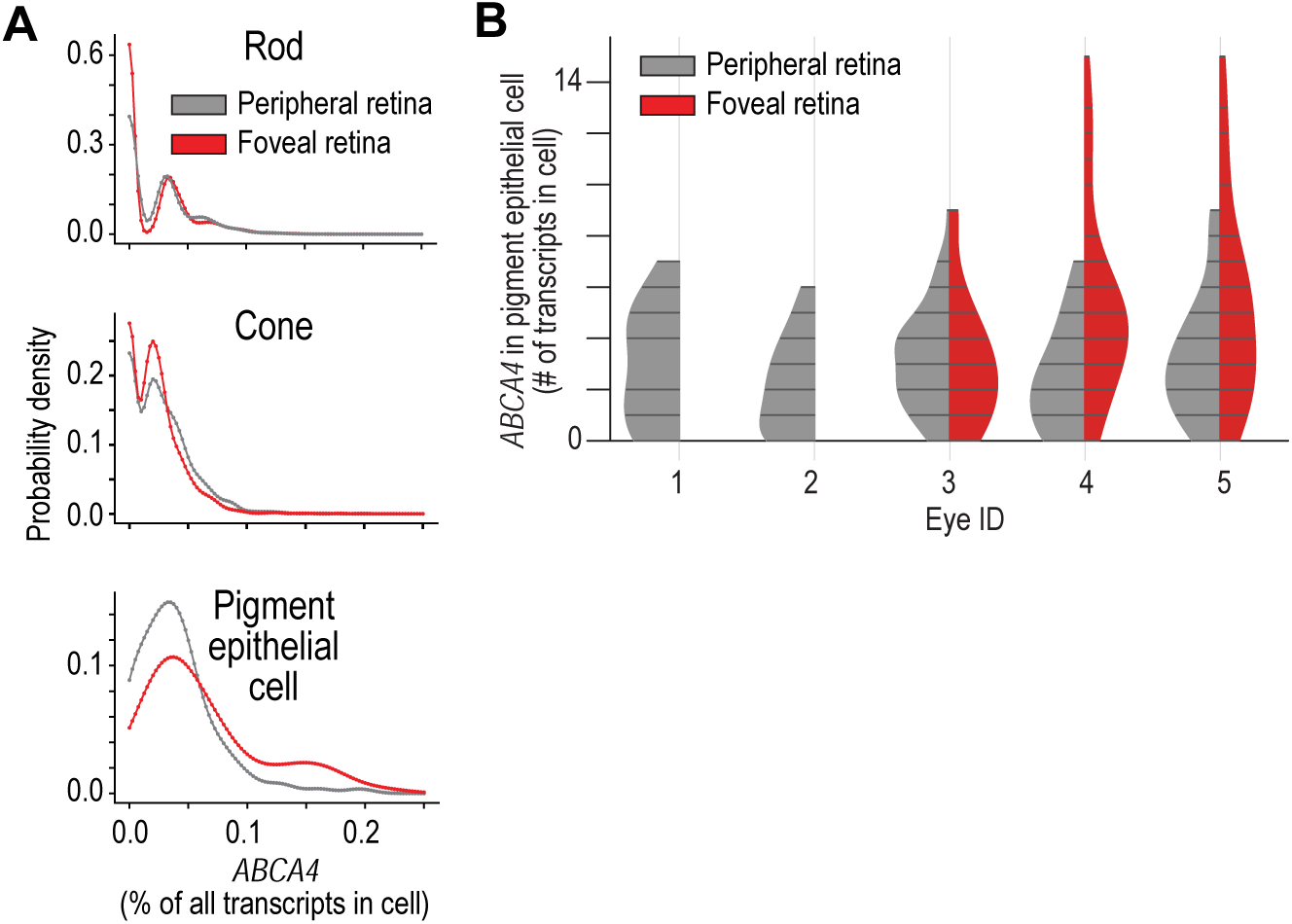
Quantification of ABCA4 expression. (A) Comparison of peripheral retina (black) and foveal retina (red) *ABCA4* expression levels in rods, cones, and pigment epithelial cells. Probability densities are generated using a Gaussian kernel density estimate with bandwidth set by Scott’s rule. (B) Violin plots show the *ABCA4* transcript counts in pigment epithelial cells. Expression is subdivided by the eye ID and region of origin. No foveal choroidal/pigment epithelial sample was available for eye IDs one and two. Eye five came from a different donor than eyes three and four.

## Supplemental Table Legend

**Supplemental Table 1.** Properties of iPSC lines tested for retinal organoid formation.

|  | Normal control (N) or disease (D) | Cell line | Formation of retinal structures on matrigel | Maintenance of layered appearance for > 100 days | Number of differentiation experiments |
| --- | --- | --- | --- | --- | --- |
| 1 | N | HPS0076: 409B2, Riken Cell Bank | no | no | 1 |
| 2 | N | iPS(IMR90)-1-DL-01, WiCell | no | no | 1 |
| 3 | N | iPS(IMR90)-4-DL-01, WiCell | yes | yes | >24 |
| 4 | N | ND41866*C, Coriell | no | no | 4 |
| 5 | N | GM23396*D, Coriell | no | no | 4 |
| 6 | N | GM23450*B, Coriell | no | no | 4 |
| 7 | N | 01F49i-N-B7 (F49B7), internal | yes | yes | >61 |
| 8 | N | Internal line 1 | yes | yes | >11 |
| 9 | D | Internal line 2 | no | no | 1 |
| 10 | N | Internal line 3 | no | no | 1 |
| 11 | N | Internal line 4 | no | no | 1 |
| 12 | D | Internal line 5 | no | no | 1 |
| 13 | D | Internal line 6 | no | no | 1 |
| 14 | N | Internal line 7 | no | no | 2 |
| 15 | D | Internal line 8 | no | no | 1 |
| 16 | D | Internal line 9 | yes | yes | 2 |
| 17 | D | Internal line 10 | yes | no | 1 |
| 18 | D | Internal line 11 | yes | yes | 1 |
| 19 | N | Internal line 12 | yes | yes | 2 |
| 20 | N | Internal line 13 | no | no | 1 |
| 21 | D | Internal line 14 | yes | no | 1 |

